# PERTURBATION OF THE AMINO ACID STRESS RESPONSE PATHWAY EXPOSES A METABOLIC VULNERABILITY IN MULTIPLE AGGRESSIVE CANCER SUBTYPES

**DOI:** 10.64898/2026.11.02.763650

**Authors:** Evelien Peeters, Igor Fijalkowski, Sara T’Sas, Kasper Thorhauge Christensen, Jordy De Coninck, Els Van Ammel, Lasse Vleminckx, Kenneth Hofkens, Lore Mennes, Alexandra Reekmans, Kaat Durinck, Kesia Van Coillie, Eleonora Leucci, Maria Francesca Baietti, Jeroen Verstappe, Panagiotis Ntziachristos, Hilde de Rooster, Filip Van Nieuwerburgh, Koen Van De Vijver, Gerben Menschaert, Tim Lammens, Pieter Van Vlierberghe, Geert Berx, Nele Loret, Maaike Van Trimpont, Steven Goossens

## Abstract

Cancer cells frequently experience nutrient limitation and depend on adaptive stress-response pathways to maintain metabolic homeostasis. Here, we show that pharmacological perturbation of amino acid stress signaling exerts broad anti-tumor activity across multiple aggressive malignancies, including acute lymphoblastic leukemia, acute myeloid leukemia, triple-negative breast cancer, ovarian cancer, and neuroblastoma. Sensitivity to pathway modulation is associated with a chronic stress-adapted metabolic state marked by constitutive amino acid response signaling, elevated NFE2L2 expression, and reduced mitochondrial abundance despite evidence of efficient oxidative metabolism. Mechanistically, we demonstrate that the ATP-competitive compound GCN2-IN-6 does not phenocopy genetic GCN2 suppression and exhibits context-dependent effects on the amino acid stress response. Nevertheless, tumors experiencing chronic metabolic stress remain exquisitely sensitive to disruption of this adaptive network, regardless of whether signaling is perturbed through canonical inhibition or paradoxical activation. Collectively, our findings identify chronic metabolic stress adaptation as an actionable vulnerability and establish off-balancing amino acid stress signaling as a therapeutic strategy across multiple cancer types.

## Introduction

Metabolic adaptation is a fundamental hallmark of cancer, enabling malignant cells to sustain proliferation in nutrient-poor microenvironments.^1,2^ Their rapid growth requires a continuous supply of nutrients and energy, creating a dependency on external sources of metabolic precursors for their macromolecule biosynthesis. These metabolic vulnerabilities have inspired therapeutic strategies aimed at nutrient deprivation, of which L-asparaginase (ASNase) therapy in acute lymphoblastic leukemia (ALL) remains the most successful clinical example.^3,4^

Cancer cells can, however, evade amino acid deprivation through adaptive stress response pathways. A key regulator of this process is the amino acid response (AAR), a branch of the integrated stress response (ISR) governed by the eIF2α kinase GCN2.^5–7^ Upon amino acid shortages, uncharged tRNAs trigger autophosphorylation of this kinase, leading to a translational rewiring with a decrease in global protein synthesis and freeing up resources for a selective translation of stress-responsive transcripts, including the upregulation of activating transcription factor 4 (ATF4). By binding C/EBP-ATF response element (CARE) sequences in promotor regions, ATF4 induces a transcriptional program that mitigates nutrient imbalances and restores metabolic homeostasis.^8^

Cancer cells hijack these stress-adaptation mechanisms to sustain their proliferative drive and to cope with their self-imposed oncogenic stress. Across multiple cancer models, GCN2 activation has been shown to promote proliferation^9–12^, metastatic progression^10,13^ and therapy resistance^14–16^. These insights have sparked interest in therapeutically targeting GCN2, leading to the development of several ATP-competitive GCN2 inhibitors.^17–20^

Here, we further evaluated the therapeutic potential of GCN2-IN-6, a previously reported selective and orally available GCN2 inhibitor^18^, in a wide series of hematological and solid cancer patient derived xenograft (PDX) models. Administration of GCN2-IN-6 exhibited robust anti-proliferative effects in most, but not all, tested models. Ribosome profiling matched with RNA sequencing revealed that responsive tumors exist in a chronically stress-adapted metabolic state characterized by constitutive activation of amino acid stress signaling, enhanced oxidative stress tolerance, and reduced mitochondrial abundance, despite evidence of preserved mitochondrial function.

The interpretation of GCN2-IN-6 as a selective inhibitor of this adaptive response is, however, complicated by emerging evidence that ATP-competitive GCN2 inhibitors can exhibit context-dependent pharmacology. Rather than uniformly suppressing GCN2 signaling, these compounds can paradoxically activate the pathway under specific conditions, raising important questions regarding their mechanism of action *in vivo*.^21–23^ Consistent with these previous reports, our results demonstrate that GCN2-IN-6 can induce either inhibition or activation of the GCN2 pathway depending on the cellular environment and fails to phenocopy genetic suppression of GCN2 *in vivo*. Nevertheless, both genetic and pharmacological studies converge on a central concept: chronically stressed tumors require precise regulation of amino acid stress signaling to maintain metabolic homeostasis. Off-balancing this finely tuned adaptive state—whether through inhibition, activation, or broader disruption of the signaling network—pushes metabolically stressed cancer cells beyond their adaptive capacity, which results in profound anti-tumor effects. Consequently, GCN2-IN-6 exerts broad anti-tumorigenic activity across multiple aggressive cancer types.

## Results

### ASNase-induced amino acid stress drives AAR-dependent adaptation and reveals a stress signaling vulnerability in ALL

For decades, asparagine deprivation therapy has been a cornerstone in the treatment of ALL.^3,4,24,25^ Leukemic blasts in ALL typically express insufficient levels of ASNS and therefore cannot synthesize their own asparagine, rendering them entirely dependent on extracellular sources of this amino acid for survival and proliferation. However, adaptive stress responses and associated metabolic rewiring can enable lymphoblasts to withstand amino acid shortages, potentially diminishing sensitivity to ASNase therapy.^17,26^ Because GCN2 is a key sensor of amino acid insufficiencies that activates ATF4-dependent adaptive stress programs, we investigated whether the administration of a GCN2-IN-6, a reported potent and orally available GCN2 inhibitor, could prevent ASNase-induced AAR pathway activation in patient-derived T-cell acute lymphoblastic leukemia (T-ALL).

To directly assess cellular metabolic adaptation under sustained asparagine deprivation, T-ALL PDX1 cells —previously shown to be sensitive to ASNase *in vivo* (Fig. S1a)— were engrafted in NOD SCID gamma (NSG) mice and treated with a single high dose (100 IU) of ASNase once leukemic burden exceeded 50% hCD45⁺ cells. Leukemic blasts were isolated five days later, during ongoing asparagine deprivation, and rechallenged *ex vivo* with increasing concentrations of ASNase. Lymphoblasts derived from ASNase-treated leukemic animals exhibited significantly reduced sensitivity to ASNase compared to vehicle-derived controls (Fig. 1a), demonstrating that leukemic cells acquire a resistant phenotype under continuous amino acid deprivation.

**FIGURE 1.**
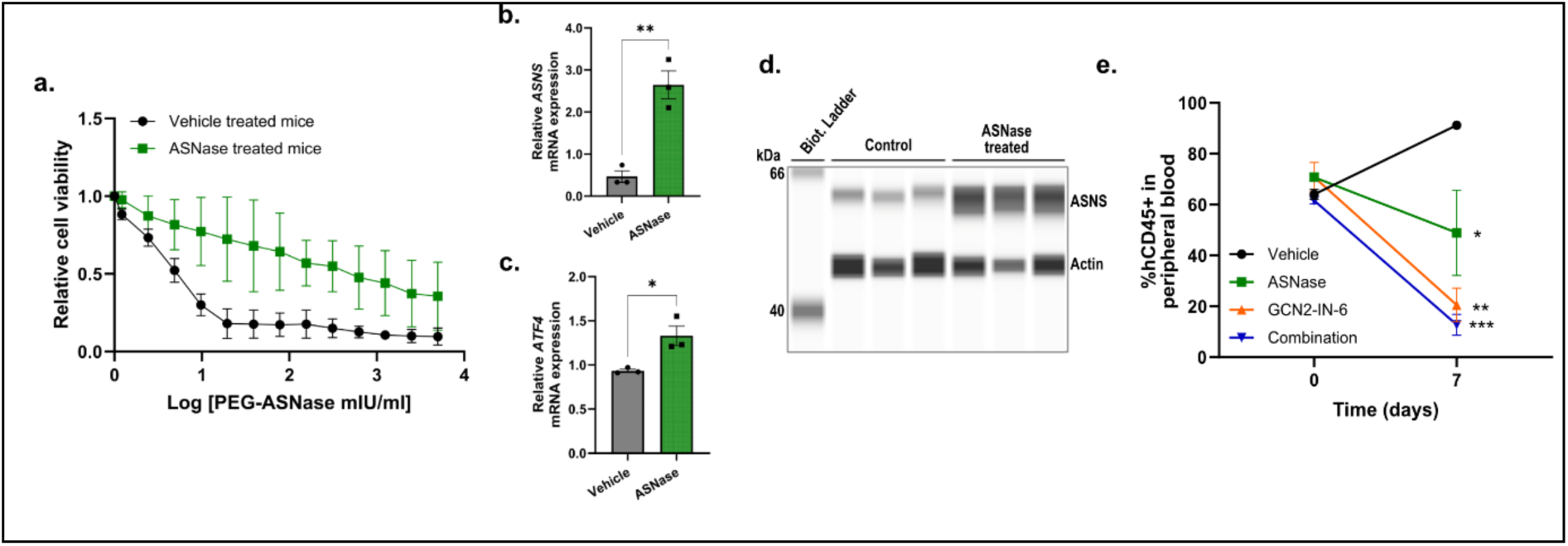
a. Relative viability of leukemic cells isolated from mice treated with a single intravenous dose of ASNase (100 IU/mouse) or vehicle (n = 3 mice per group). Cells were treated *ex vivo* with a two-fold serial dilution of ASNase (maximum concentration 2000 mIU/mL) for 72 h before viability was assessed using CellTiter-Glo. Concentrations were log-transformed. Each dot represents the mean of three biological replicates; error bars indicate SD. b. Relative *ASNS* mRNA expression in leukemic cells isolated from mice in (a) treated with ASNase or vehicle. Error bars indicate SEM. **p = 0.0038 (unpaired two-tailed t-test). c. Relative *ATF4* mRNA expression in leukemic cells isolated from mice in (a) treated with ASNase or vehicle. Error bars indicate SEM. *p = 0.0238 (unpaired two-tailed t-test). d. Automated capillary western blot (WES) showing ASNS protein expression in leukemic cells isolated from mice in *a* treated with ASNase or vehicle. β-Actin serves as the loading control. Each lane is a biological replicate. e. Leukemic burden in T-ALL PDX1 mice, measured as the percentage of human CD45-positive (hCD45⁺) cells in peripheral blood. Mice received vehicle, ASNase (25 IU/mouse, single dose; n = 3), GCN2-IN-6 (3 mg/kg, oral dose every 3.5 days; n = 3), or the combination (n = 3). No significant differences were observed between treatment groups before treatment (day 0). At day 7, all treatment groups showed reduced leukemic burden compared with vehicle (*p = 0.0300 for ASNase, **p = 0.0017 for GCN2-IN-6, ***p = 0.0009 for the combination; one-way ANOVA with Dunnett’s multiple-comparison correction). Dots represent group means; error bars indicate SEM.

Mechanistically, acquisition of this tolerant phenotype coincided with activation of the AAR pathway. Leukemic cells isolated from ASNase-treated mice displayed increased expression of *ATF4* (*p = 0.0238; unpaired t-test) and its downstream target asparagine synthetase (*ASNS)* (**p=0.0038; unpaired t-test) at the mRNA level, accompanied by increased ASNS protein abundance (Fig. 1b-d). Collectively, these data demonstrate that sustained ASNase exposure drives AAR pathway activation in ALL to overcome the inflicted metabolic stress and tolerate continued systemic asparagine depletion.

We next questioned whether disrupting this adaptive stress response could enhance the efficacy of ASNase therapy. Mice bearing advanced leukemia (>50% hCD45⁺) were randomized to receive ASNase (25 IU/mouse, 1 injection IP, n = 3), GCN2-IN-6 (3 mg/kg, oral dose every 3.5 days, n = 3), the combination (n=3) or vehicle as control (n=3). Combination therapy resulted in rapid enhanced anti-leukemic effect compared to ASNase alone (Fig. 1e; ***p(combination) = 0.0009), *p(ASNase) = 0.0300, one-way ANOVA), consistent with disruption of stress-induced metabolic adaptation. However, this increased efficacy was accompanied by greater toxicity, reflected by enhanced body weight loss (Fig. S1b), suggesting that simultaneous targeting of amino acid bioavailability and the stress-response signaling narrows the therapeutic window. Unexpectedly, GCN2-IN-6 monotherapy exerted anti-leukemic effects that exceeded those of ASNase (Fig. 1e; **p(GCN2-IN-6) = 0.0017, one-way ANOVA). Moreover, the enhanced efficacy observed with the combination treatment appeared largely attributed to GCN2-IN-6. This finding contrasts with prior *in vitro* studies under optimal growth conditions, where GCN2 inhibition primarily sensitized cells to amino acid deprivation rather than directly impairing proliferation.^17^

### Triple negative breast cancer PDXs exhibit an *in vivo* vulnerability to perturbation of amino acid stress signaling

Given their spatially heterogeneous nutrient availability, we hypothesized that solid cancers may be particularly dependent on amino acid stress response signaling. In breast cancer, and particularly in triple-negative breast cancer (TNBC), metabolic rewiring is a defining feature that supports tumor growth, plasticity, and metastatic dissemination.^27–30^ These tumors frequently encounter fluctuating nutrient availability *in vivo*, suggesting a potential reliance on adaptive stress-response pathways to maintain metabolic homeostasis.^26^ We therefore investigated whether TNBC cells depend on AAR signaling and whether perturbation of this pathway represents a therapeutic vulnerability.

To functionally assess this dependency, we evaluated the response of the ASNS-low MDAMB231 cells to amino acid stress *in vitro*. Under standard culture conditions, pharmacological targeting of GCN2 using GCN2-IN-6 had no measurable effect on cell proliferation (IC_50_ not reached), consistent with the absence of nutrient limitation and minimal activation of stress-response pathways (Fig. 2a). Induction of amino acid stress using ASNase resulted in a modest growth inhibition (IC_50_ = 1.42 IU/mL), accompanied by activation of the AAR pathway, as evidenced by increased phosphorylated GCN2 (pGCN2) and induced ASNS expression (Fig. 2b). Notably, combined treatment with ASNase and GCN2-IN-6 markedly enhanced growth inhibition (Fig. 2a; IC_50_ = 0.009 IU/mL), indicating that perturbation of stress-response signaling sensitizes TNBC cells to amino acid deprivation. This went paired with an inhibition of the GCN2-ATF4 pathway.

**FIGURE 2.**
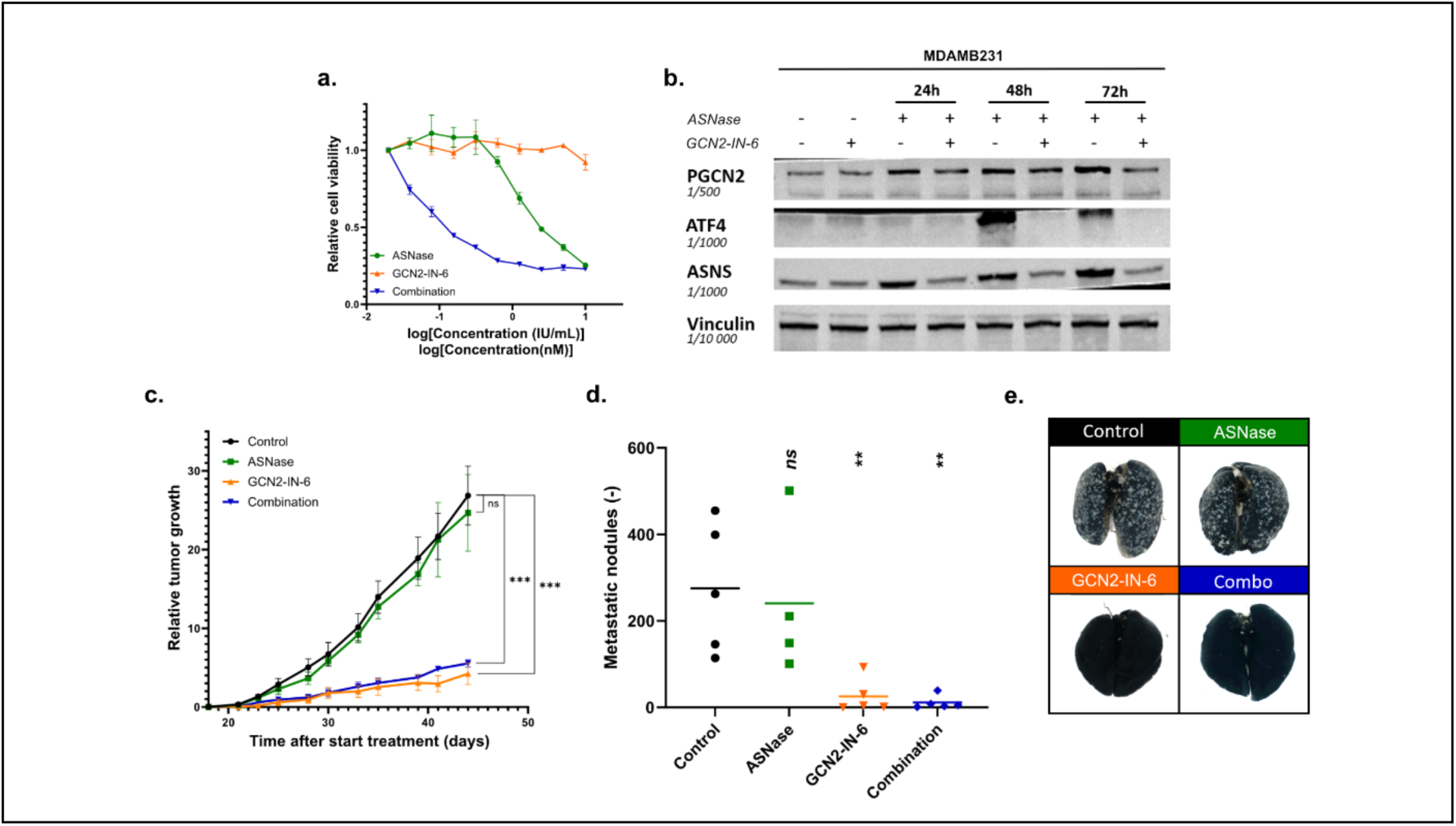
**a.** Relative cell viability of MDAMB231 cells treated with ASNase (maximum concentration 10 IU/mL), GCN2-IN-6 (maximum concentration 2000 nM) or the combination for 72 h. Both compounds were administered as two-fold serial dilutions. Viability was assessed using CellTiter-Glo. IC₅₀ values were determined by nonlinear regression: IC_50_(ASNase) = 1.42 IU/mL, IC_50_(GCN2-IN-6) could not be determined, IC_50_(combination) = 0.009 IU/mL. **b.** Representative western blot analysis of phosphorylated GCN2 (pGCN2), ATF4, and ASNS protein expression in MDAMB231 cells treated with ASNase (1 IU/mL), GCN2-IN-6 (200 nM), or the combination for 24, 48, or 72 h. Vinculin serves as the loading control. **c.** Tumor growth in orthotopic MDAMB231 xenografts following treatment with vehicle, ASNase (25 IU/mouse, intraperitoneally every 8 days), GCN2-IN-6 (3 mg/kg, orally every 4 days), or the combination (n = 5 mice per group). Treatment was initiated when tumors became palpable. Dots represent group means; error bars indicate SEM. Statistical analysis was performed using a linear mixed-effects model (REML) with time and treatment group as fixed effects. Treatment groups were compared with the vehicle control. ***p < 0.001; ns = 0.222. **d.** Ǫuantification of pulmonary metastatic nodules in mice shown in **(c)**. Metastases were visualized manually after injection with Indian ink **(see e)**. Dots represent individual mice; error bars indicate SEM. One-way ANOVA with Dunnett’s multiple-comparison test was used to compare each treatment group with vehicle: **p = 0.0093 for GCN2-IN-6, **p = 0.0062 for the combination, ns = 0.9416 for PEG-ASNase. **e.** Representative images of Indian ink-injected lungs of mice from **(d)**. Metastatic nodules appear as white lesions against the black-stained lung tissue.

Therapeutic relevance of these findings were further assessed *in vivo* using an orthotopic MDAMB231 xenograft model in NSG mice. Tumor bearing mice were randomized into four treatment groups to receive vehicle only (n = 5), ASNase (25 IU/mouse administered IP every 8 days, n = 5), GCN2-IN-6 (3 mg/kg administered orally every 4 days, n = 5), or the combination (n = 5). ASNase monotherapy did not significantly impair TNBC primary tumor growth (Fig. 2c; ns(ASNase) = 0.222, REML analysis) and no significant influence of ASNase was observed on the metastatic burden in this model (Fig. 2d-e, Fig. S2a; ns(ASNase) = 0.9416, one-way ANOVA), although variability in spontaneous metastatic dissemination limited statistical power. In contrast, GCN2-IN-6 administration resulted in a pronounced reduction in primary tumor growth, both as monotherapy (***p < 0.001, REML analysis) and in combination with ASNase (***p < 0.001, REML analysis) (Fig. 2c). Notably, GCN2-IN-6 monotherapy achieved efficacy comparable to the combination treatment, indicating that monotherapy alone is sufficient to impair tumor growth *in vivo*. In addition, GCN2-IN-6 significantly reduced metastatic burden in the lungs (**p(GCN2-IN-6) = 0.0093, one-way ANOVA) and other organs (Fig. 2d-e, Fig. S2a, Table S1), although it remains unclear whether this reflects a direct effect on metastatic capacity or is secondary to reduced primary tumor growth.

Combined treatment with ASNase did not enhance anti-tumor effects and led to increased toxicity, as evidenced by significant body weight loss (Fig. S2b), suggesting a limited therapeutic advantage for combination strategies in solid tumors. Based on these findings, we focused in subsequent analyses on GCN2-IN-6 monotherapy to further define determinants of response.

The discrepancy between *in vitro* and *in vivo* responses to GCN2-IN-6 monotherapy suggests that cancer cells within the *in vivo* tumor microenvironment may experience environmental metabolic stress, resulting in constitutive basal activation of AAR/ISR signaling pathways. Under these conditions, GCN2-IN-6 may disrupts this adaptive state, leading to impaired tumor growth even in the absence of additive amino acid deprivation therapy. An alternative explanation is that GCN2-IN-6 may engage distinct modes of action *in vitro* versus *in vivo*.

### Addressing GCN2-IN-6 specificity reveals context-dependent activation of AAR pathway

Pharmacological targeting of GCN2 has recently been complicated by reports questioning the specificity and mode of action of ATP-competitive inhibitors, including GCN2-IN-6.^21,23^ To determine whether the anti-tumor effects observed in our *in vivo* models are mediated through canonical inhibition of GCN2, we compared pharmacological inhibition with genetic suppression of GCN2. Reliable epigenetic silencing of GCN2 was achieved using CRISPR interference (CRISPRi) in MDAMB231 cells, resulting in an efficient knockdown at both the mRNA and protein levels (Fig. S3a–b). Because functional redundancy has been described between GCN2 and PERK within the integrated stress response, we also assessed PERK expression, which remained unchanged upon GCN2 knockdown (Fig. S3c).

Strikingly, genetic suppression of GCN2 failed to phenocopy the effects of GCN2-IN-6. *In vitro*, efficient GCN2 knockdown did not enhance sensitivity to ASNase (Fig. 3a), in contrast to the pronounced synergistic effects observed upon pharmacological treatment. Similarly, *in vivo*, GCN2 knockdown did not significantly affect tumor growth of MDAMB231 xenografts (Fig. 3b; Fig. S3d-e), despite confirmed knockdown in tumor tissue at endpoint (Fig. S3f). These finding demonstrate that the anti-tumor activity of GCN2-IN-6 cannot be explained solely by loss of GCN2 function.

**FIGURE 3.**
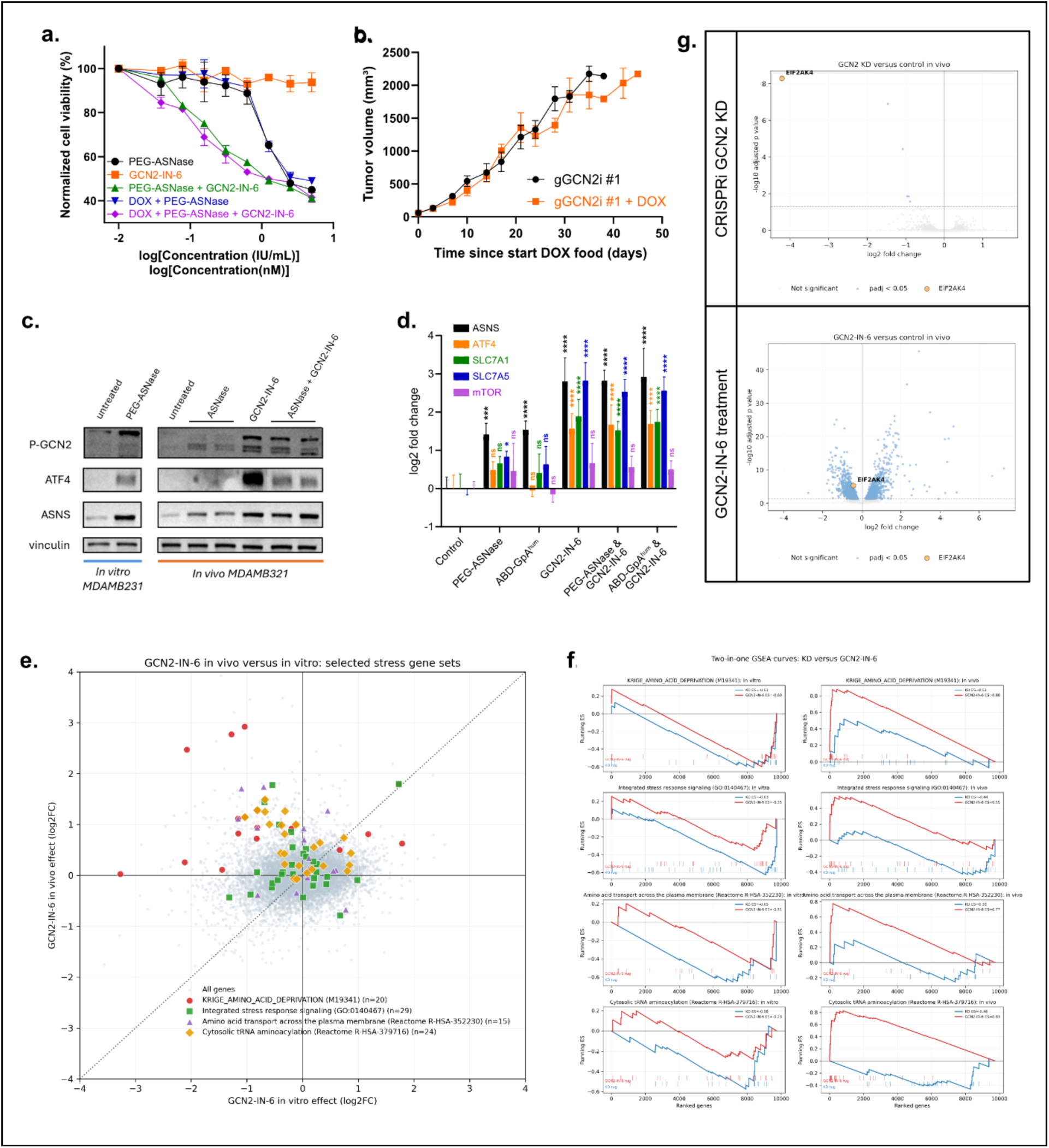
**a.** Relative cell viability of MDAMB231 gGCN2i #1 cells with or without doxycycline-induced GCN2 knockdown (1 μg/mL), treated with ASNase (maximum concentration 10 IU/mL), GCN2-IN-6 (maximum concentration 2000 nM) or the combination for 72 h. Both compounds were administered as two-fold serial dilutions. Viability was measured using CellTiter-Glo. IC₅₀ values were determined by nonlinear regression: IC_50_(PEG-ASNase) = 1.177 IU/mL, IC_50_(GCN2-IN-6) could not be determined, IC_50_(PEG-ASNase + GCN2-IN-6) = 0.2134 IU/mL, IC_50_(DOX + PEG-ASNase) = 1.143 IU/mL, IC_50_(DOX + PEG-ASNase + GCN2-IN-6) = 0.1574 IU/mL. **b.** Tumor volume of orthotopic MDAMB231 gGCN2i #1 xenografts with or without doxycycline-induced GCN2 knockdown. Doxycycline was administered continuously in the diet (200 ppm). Dots represent group means (n = 5 mice per group); error bars indicate SEM. **c.** Western blot analysis of pGCN2, ATF4, and ASNS protein expression in MDAMB231 cells treated *in vitro* with vehicle or ASNase and in tumors harvested from mice treated with vehicle, ASNase, GCN2-IN-6, or the combination. Vinculin serves as the loading control. **d.** Relative mRNA expression levels (log2 fold change) of *ASNS, ATF4, SLC7A1, SLC7A5* and *mTOR* in tumors from mice treated with vehicle, ASNase (25 IU/mouse every 8 days), GCN2-IN-6 (3 mg/kg twice weekly), or the combination. Statistical significance was determined by one-way ANOVA relative to the vehicle-treated group. Significance is indicated in the figure (ns > 0.05, * < 0.05, ** <0.01, *** <0.001, ****<0.0001). **e.** Scatter plot of the GCN2-IN-6 in vivo effect (GCN2-IN-6 treatment versus control in vivo, log2FC) versus GCN2-IN-6 in vitro effect (GCN2-IN-6 treatment versus control in vitro, log2FC). Gene sets related to the GCN2 stress response are highlighted (KRIGE_AMINO_ACID_DEPRIVATION (MSigDB identifier M19341), Integrated stress response signaling (Gene Ontology GO:0140467), Amino-acid transport across the plasma membrane (Reactome R-HSA-352230), and Cytosolic tRNA aminoacylation (Reactome R-HSA-37916). **f.** Gene set enrichment analysis (GSEA) of gene sets associated with amino acid stress signaling following CRISPRi-mediated GCN2 knockdown (blue) or GCN2-IN-6 treatment (red) under *in vitro* and *in vivo* conditions. **g.** Volcano plots of differentially expressed genes identified by bulk RNA sequencing following CRISPRi-mediated GCN2 knockdown or GCN2-IN-6 treatment under *in vivo* conditions. Differentially expressed genes (adjusted *p* < 0.05) are shown in blue, and *EIF2AK4 (GCN2)* is highlighted in orange.

Recent studies have proposed that ATP-competitive GCN2 inhibitors may exhibit biphasic behavior, functioning as inhibitors at saturating concentrations but paradoxically promoting GCN2 activation under sub-saturating conditions through stabilization of an active kinase conformation.^21,23^ This concentration-dependent phenomenon was demonstrated *in vitro* by Carlson et al.^21^ Consistent with this model, analysis of ISR pathway activation *in vivo* revealed that GCN2-IN-6 treatment induces, rather than suppresses, GCN2 signaling. Protein analysis of MDAMB231 tumors harvested after prolonged *in vivo* treatment demonstrated that GCN2-IN-6, either as monotherapy or in combination with L-asparaginases, resulted in marked upregulation of phosphorylated GCN2 (pGCN2), ATF4, and ASNS (Fig. 3c). In contrast, ASNase treatment alone induced ASNS expression with only modest increases in pGCN2 and minimal changes in ATF4 protein levels. These observations suggest that GCN2-IN-6 promotes activation of the GCN2–ATF4 axis *in vivo*, rather than inhibition. This effect was validated at the transcriptional level (Fig. 3d). GCN2-IN-6 treatment significantly increased expression of canonical ISR target genes, including *ATF4*, *ASNS*, *SLC7A1*, and *SLC7A5*, exceeding the induction observed with L-asparaginase treatment alone.

To determine whether this paradoxical activation is context-dependent, we performed bulk RNA sequencing comparing GCN2-IN-6-treated MDAMB231 cells *in vitro* and *in vivo*. Strikingly, gene sets associated with amino acid deprivation and ISR activation were upregulated *in vivo* but downregulated *in vitro* following GCN2-IN-6 treatment (Fig. 3e), validating paradoxical activation *in vivo*.

Of interest, GCN2-IN-6 induced markedly more transcriptional changes than GCN2 knockdown (Fig. 3g). Other pathways downregulated upon GCN2-IN-6 treatment *in vivo* were associated with epithelial to mesenchymal transition (EMT), and extracellular matrix remodeling (Fig. S3g-i). The drug appears to reproducibly suppress a specific transcriptional state associated with mesenchymal identity, tumor plasticity and invasive behavior. Additionally, an environment-independent reproducible reduction of expression of SUZ12 and SOX2 was observed after GCN2-IN-6 treatment, two known regulators of tumorigenesis and metastasis.^31–34^ This suggests that the transcriptional effects observed with GCN2-IN-6 in TNBC may not exclusively be mediated through on-target GCN2 inhibition and may involve a broader indirect stress-response, secondary cellular responses or off-target kinase effects. This mechanistic framework provides an explanation for the discrepancy between genetic and pharmacological perturbation of GCN2 and highlights the importance of cellular context in determining the functional outcome of GCN2-targeting strategies.

### Anti-cancer effects of GCN2-IN-6 monotherapy is reproducible in multiple tumor types

To assess the generalizability of our findings, we evaluated the therapeutic efficacy of GCN2-IN-6 across a broader panel of patient-derived xenograft (PDX) models representing both hematological and solid malignancies. In all cases, GCN2-IN-6 was administered orally at a dose of 3 mg/kg twice per week, and was compared against vehicle-treated mice.

A diverse set of T-ALL and B-ALL PDX models was selected, each exhibiting distinct baseline expression profiles of AAR pathway components (Fig. S4a-b). Despite this heterogeneity, GCN2-IN-6 treatment consistently suppressed leukemic progression in all models, both of T-ALL and B-ALL origin (Fig. 4a–f, statistics in figure), indicating that *in vivo* sensitivity to GCN2-IN-6 is broadly conserved across lymphoid malignancies. Notably, baseline *ATF4* expression did not predict therapeutic response, as all ALL PDXs responded to GCN2-IN-6 despite variable *ATF4* expression levels. Treatment was well tolerated, with only mild and reversible effects on body weight (Fig. S4c–h). In all models, tumor growth resumed upon discontinuation of GCN2-IN-6, indicating that sustained treatment is required to maintain disease control.

**FIGURE 4.**
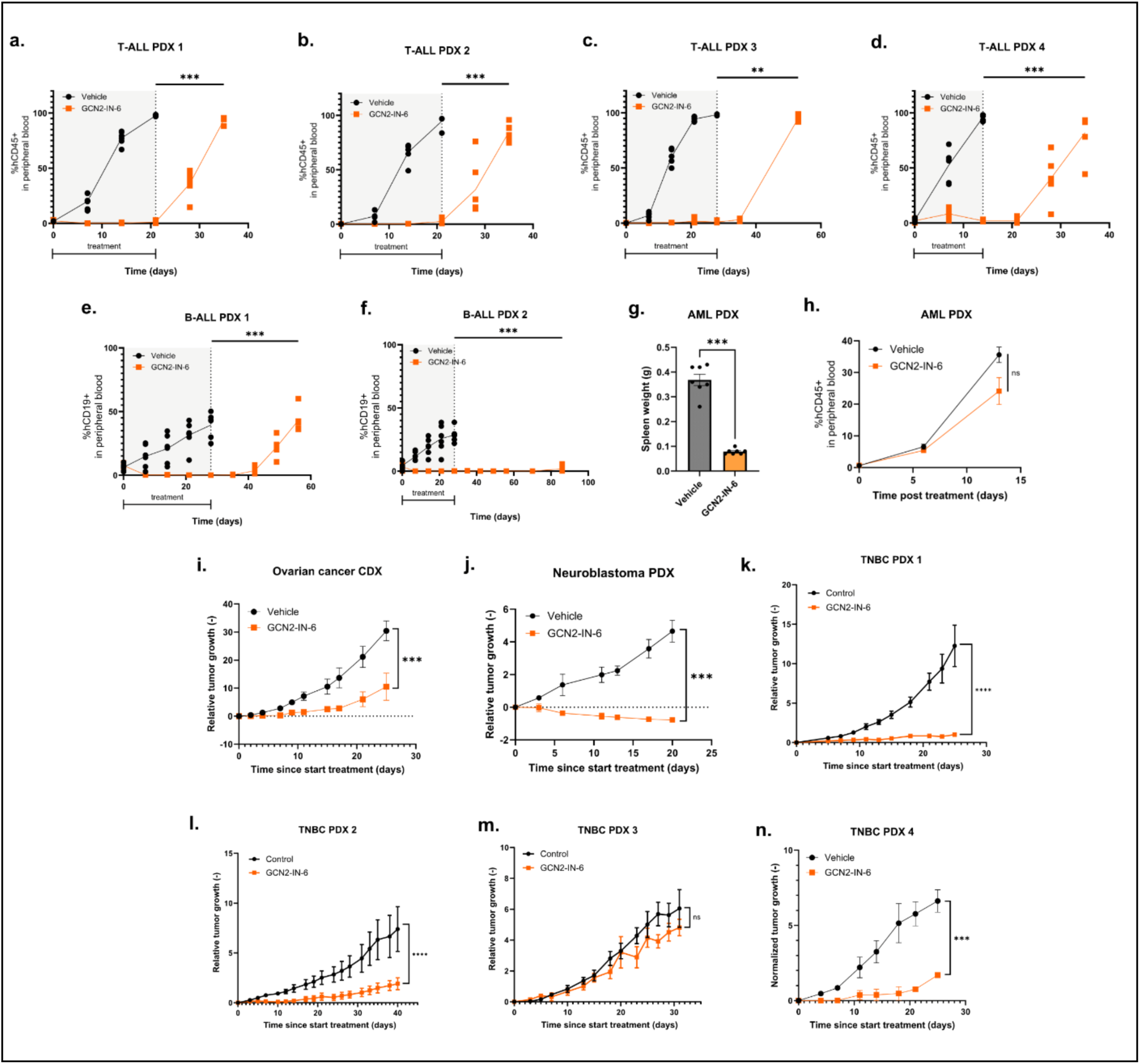
**a-d.** Leukemic burden, measured as the percentage of human CD45-positive (hCD45⁺) cells in peripheral blood, in T-ALL PDX models 1–4 treated with vehicle or GCN2-IN-6 (3 mg/kg, orally, twice weekly; n = 6 mice per group). Treatment was discontinued once vehicle-treated mice reached the experimental endpoint, and the remaining mice were monitored for survival. Survival was analyzed using the Kaplan-Meier method with log-rank (Mantel-Cox) testing. Significance is indicated in the figure. **e-f.** Leukemic burden, measured as the percentage of human CD19-positive (hCD19⁺) cells in peripheral blood, in B-ALL PDX models 1 and 2 treated with vehicle or GCN2-IN-6 (3 mg/kg, orally, twice weekly; n = 6 mice per group). Treatment was discontinued once vehicle-treated mice reached the experimental endpoint, and the remaining mice were monitored for survival. Survival was analyzed using the Kaplan-Meier method with log-rank (Mantel-Cox) testing. Significance is indicated in the figure. **g.** Spleen weight of AML PDX-bearing mice treated with vehicle or GCN2-IN-6 (3 mg/kg, orally, twice weekly; n = 7 mice per group). Spleens were collected at the experimental endpoint. Statistical significance was determined using the Kruskal-Wallis test. **p = 0.0012. **h.** Leukemic burden, measured as the percentage of hCD45⁺ cells in peripheral blood, in the AML PDX model following treatment with vehicle or GCN2-IN-6 (3 mg/kg, orally, twice weekly; n = 7 mice per group). Statistical analysis was performed using a linear mixed-effects model (REML) with time and treatment as fixed effects. *p = 0.0325. **i.** Relative tumor growth of an ovarian cancer cell line-derived xenograft (CDX) treated with vehicle or GCN2-IN-6 (3 mg/kg, orally, twice weekly; n = 6 mice per group). Statistical analysis was performed using a linear mixed-effects model (REML). ***p = 0.0008 **j.** Tumor volume of a neuroblastoma PDX treated with vehicle or GCN2-IN-6 (3 mg/kg, orally, twice weekly; n = 4 per group). Statistical analysis was performed using a linear mixed-effects model (REML). ***p = 0.0002. **k – n.** Relative tumor growth of four TNBC PDX models treated with vehicle or GCN2-IN-6 (3 mg/kg, orally, twice weekly; n = 5 mice per group). Dots represent group means; error bars indicate SEM. Statistical analysis was performed using a linear mixed-effects model (REML). Significance is indicated in the figure.

We extended our analysis to acute myeloid leukemia (AML), a myeloid malignancy with distinct biology but similarly profound metabolic demands. Although therapeutic effects of GCN2-IN-6 in this single aggressive AML PDX model were less pronounced compare to ALL, GCN2-IN-6 treatment still produced a significant reduction in leukemic burden as reflected by the decreased splenomegaly (Fig. 4g, **p = 0.0012, Mann-Whitney test). However, %hCD45 in peripheral blood was not significantly different (Fig. 4h, Fig. S4i, p = 0.0325, mixed-effects model). Interestingly, AML cells treated with GCN2-IN-6 also showed activation of the AAR pathway (Fig. S4j-n) As acute leukemias are highly heterogeneous malignancies driven by a large variety of genetic perturbations, more research will be necessary to evaluate whether certain genetic subclasses are more sensitive to GCN2-IN-6 than others.

Similarly, also in most tested solid cancer xenograft models, a clear therapeutic effect was seen upon GCN2-IN-6 administration. Anti-tumor activity was observed in an ovarian cancer cell line–derived xenograft (CDX) (Fig. 4i), a neuroblastoma PDX model (Fig. 4j) and multiple triple negative breast cancer PDX models (TNBC) (Fig. 4k-n). Notably, in the neuroblastoma model, treatment resulted in tumor regression, indicating a particularly strong dependency on balanced stress response signaling in this context. In all models, treatment was well tolerated (Fig. S4o-t).

Of the TNBC models, sensitivity varied with three sensitive and one resistant model. These results contrast with the more uniform sensitivity observed in ALL. This variability prompted us to investigate the molecular determinants of response, with a particular focus on TNBC, to identify potential biomarkers that can stratify sensitive and resistant tumors.

### A chronic stress-adapted metabolic state predicts sensitivity to GCN2 pathway perturbation

To identify molecular determinants underlying differential sensitivity to GCN2-IN-6, we performed matched ribosome profiling and RNA sequencing on untreated primary tumors from the most sensitive (TNBC PDX 1) and resistant (TNBC PDX 3) models (3 mice/model) (*Fig. S5a-h)*. Transcriptional and translational data were integrated using the DeltaTE framework^35^ (Fig. 5a), enabling discrimination between transcriptionally regulated genes (DTG; Fig. S5a) and genes whose expression is primarily controlled at the level of translation (DTEG; Fig. S5b).

**FIGURE 5.**
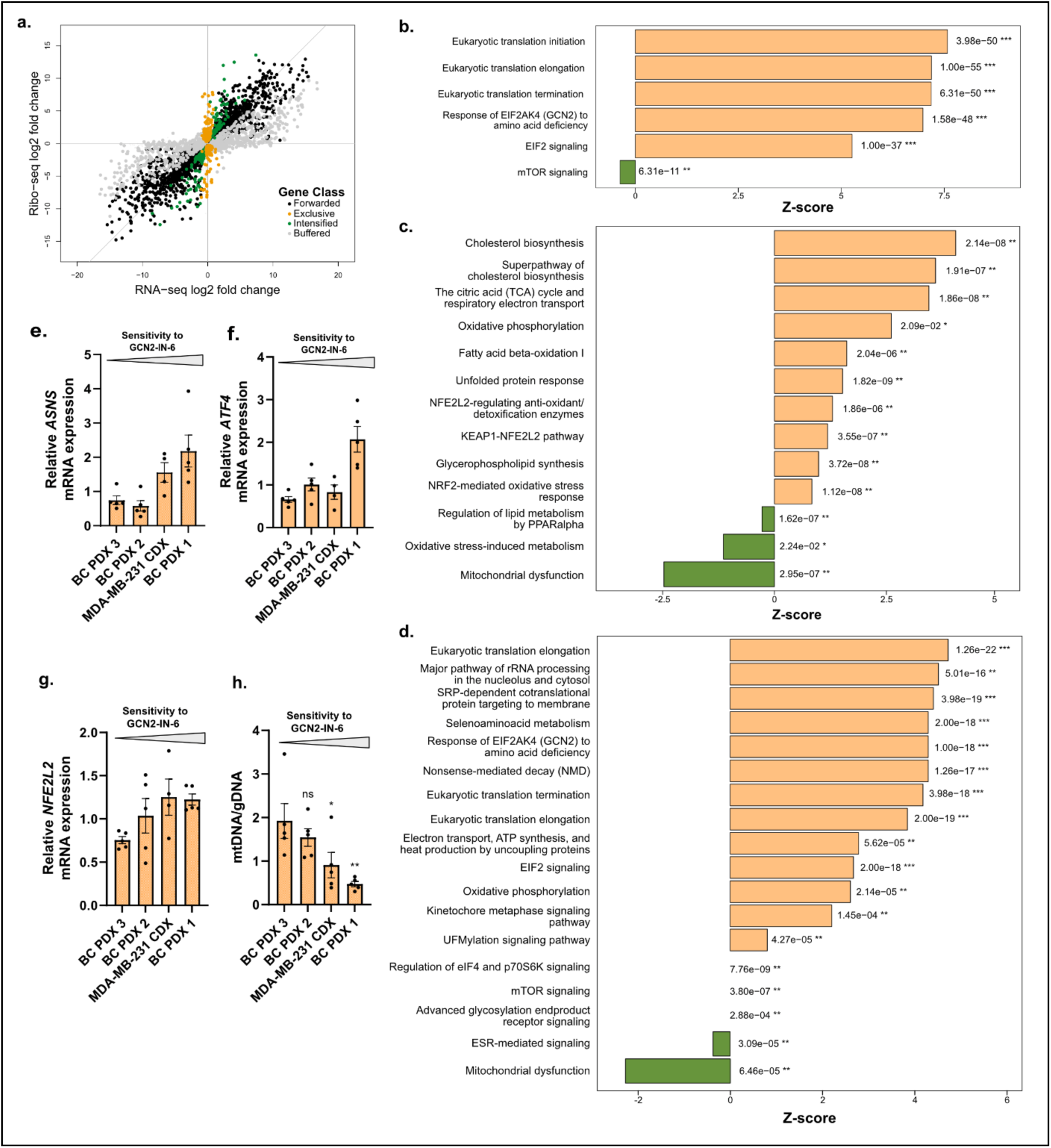
**a.** Scatter plot comparing differential gene expression measured by bulk RNA sequencing and ribosome profiling (RIBOseq). Genes were classified according to the DeltaTE method into forwarded, exclusive, intensified or buffered. Each dot represents a gene. **b.** Pathway enrichment analysis of genes belonging to the buffered cluster, comparing TNBC PDX 1 with TNBC PDX 3 as the reference. Selected pathways associated with the amino acid response (AAR) and protein translation are shown. Bars represent activation *z*-scores and are ordered by decreasing *z*-score. Positive *z*-scores indicate predicted activation in TNBC PDX 1 relative to TNBC PDX 3, whereas negative *z*-scores indicate predicted inhibition. Associated p values and significance levels are indicated next to each bar. **c.** Pathway enrichment analysis comparing TNBC PDX 1 with TNBC PDX 3 as the reference. Selected differentially regulated pathways associated with cellular stress responses, oxidative phosphorylation, and mitochondrial function are shown. Bars are ordered by decreasing *z*-score. Positive *z*-scores indicate predicted pathway activation in TNBC PDX 1 relative to TNBC PDX 3, whereas negative *z*-scores indicate predicted inhibition. Associated p values and significance levels are indicated next to each bar. **d.** Pathway enrichment analysis of bulk RNA-seq data from patients with ATF4-high tumors compared with the remaining patients in the METABRIC breast cancer cohort. Selected differentially regulated pathways are shown. Bars are ordered by decreasing *z*-score. Positive and negative *z*-scores indicate predicted pathway activation and inhibition, respectively, in the ATF4-high cohort. Associated p values and significance levels are indicated next to each bar. **e-h.** Baseline expression of candidate biomarkers across breast cancer models with increasing sensitivity to GCN2-IN-6 (indicated by the gradient above each graph). Relative mRNA expression of **(e)** ASNS, **(f)** ATF4, and **(g)** NFE2L2, and **(h)** relative mitochondrial DNA content normalized to genomic DNA (mtDNA/gDNA) were determined by RT-qPCR in BC PDX3, BC PDX2, MDAMB231 CDX, and BC PDX1. Bars represent mean ± SEM, with each dot representing an individual biological replicate. Statistical significance is indicated in the figure. Statistical analysis was performed using one-way ANOVA.

We first questioned whether sensitive tumors exhibit evidence of constitutive amino acid stress. Analysis of genes displaying buffered translation—where transcriptional induction is not accompanied by increased translation efficiency, consistent with translational restraint during amino acid limitation—revealed marked enrichment of amino acid response pathways in the sensitive model. Gene sets associated with response of EIF2AK4 (GCN2) to amino acid deficiency (Z=7.076, ***p=1.58e-48) and EIF2 signaling (Z=5.333, ***p=1.00e-37) were among the most significantly enriched pathways (Fig. 5b), demonstrating that GCN2-IN-6 sensitive tumors exist in a state of chronic amino acid stress before therapeutic intervention. Of interest, codon occupancy analysis revealed the amino acid stress could not be attributed to a specific amino acid deficiency (Fig. S5i) and pathways involved in translation initiation, elongation and termination were among the strongest positively enriched signatures (Fig. 5b). This indicates a state of chronic amino acid stress in which translational capacity is continuously remodeled to sustain protein synthesis despite persistent nutrient limitation.

We next investigated how sensitive tumors adapt to this chronic metabolic challenge. Analysis of pathways with increased translation efficiency demonstrated coordinated activation of multiple stress adaptation programs rather than isolated pathway alterations. Sensitive tumors displayed enrichment of the unfolded protein response together with robust activation of NRF2-mediated oxidative stress pathways, including the KEAP1–NFE2L2 axis and antioxidant detoxification programs (Fig. 5c). These findings indicate that responsive tumors experience persistent oxidative stress and rely on constitutive activation of cytoprotective mechanisms to maintain redox homeostasis.

Remarkably, these stress adaptation programs were accompanied by extensive metabolic rewiring centered on mitochondrial function. Sensitive tumors exhibited increased tricarboxylic acid (TCA) cycle activity, oxidative phosphorylation, fatty acid β-oxidation, glycerophospholipid synthesis and cholesterol biosynthesis, while pathways associated with mitochondrial dysfunction were significantly depleted (Fig. 5c). Rather than reflecting mitochondrial impairment, this pattern is consistent with highly efficient mitochondrial metabolism supporting continued proliferation under conditions of chronic nutrient stress. Increased lipid and cholesterol biosynthesis further suggest that mitochondrial-derived metabolites are actively redirected toward membrane production and biomass accumulation, despite ongoing amino acid limitation.

To determine whether these observations extend beyond our experimental models, we interrogated publicly available breast cancer METABRIC transcriptomic dataset. Tumors exhibiting constitutive activation of amino acid stress signaling similarly displayed enrichment of oxidative phosphorylation and reduced mitochondrial dysfunction (Fig. 5d), indicating that this metabolic phenotype is conserved in human breast cancer.

We next evaluated whether components of this stress-adapted metabolic program correlate with therapeutic response across the complete TNBC xenograft panel. Among the candidate genes examined, expression of the canonical AAR components *ATF4* and *ASNS* showed a positive association with sensitivity to GCN2-IN-6, with the highest expression observed in the most responsive models (Fig. 5e–f). Similarly, expression of NFE2L2, a central regulator of cellular redox homeostasis and mitochondrial adaptation, progressively increased with increasing sensitivity to GCN2-IN-6 (Fig. 5g). Because transcriptional profiling suggested altered mitochondrial function, we quantified mitochondrial abundance by determining the mitochondrial DNA (mtDNA) to genomic DNA (gDNA) ratio. Strikingly, mitochondrial content progressively decreased with increasing sensitivity to GCN2-IN-6 (Fig. 5h).

Together with the observed increase in oxidative phosphorylation and depletion of mitochondrial dysfunction pathways, these findings suggest that sensitive tumors possess fewer but metabolically more efficient mitochondria. We therefore propose that these tumors exist in a highly adapted metabolic state characterized by constitutive amino acid stress signaling, efficient oxidative metabolism, enhanced lipid biosynthesis and robust antioxidant defenses. While this coordinated adaptation enables sustained tumor growth under nutrient-limited conditions, it simultaneously creates a dependence on adaptive stress-response signaling, rendering these tumors selectively vulnerable to pharmacological perturbation of the GCN2 pathway.

## Discussion

Adaptive stress responses allow cancer cells to survive fluctuating nutrient conditions and represent an underexplored therapeutic vulnerability. In the present study, we demonstrate that perturbation of amino acid stress signaling exerts broad anti-tumor activity across multiple aggressive malignancies, including T-ALL, B-ALL, AML, TNBC, neuroblastoma and ovarian cancer. Importantly, our findings indicate that tumor sensitivity is determined not by a specific amino acid dependency, but rather by the extent to which tumors rely on constitutive stress adaptation programs to maintain metabolic homeostasis.

ALL patients are currently treated with an intensified combination of chemotherapeutics and ASNases.^36,37^ The inclusion of ASNase therapy has drastically improved overall outcome of pediatric ALL patients.^37–39^ However, administration of the clinically approved bacterial derived ASNases remains suboptimal as it is accompanied by numerous immunogenic and non-immunogenic side-effects^40^. These tolerability issues often necessitate exclusion of ASNases from the treatment regimen, prompting the search for novel approaches targeting this amino acid dependency. Our data indicate that GCN2 inhibition therapy could serve as an alternative for ASNases as part of first- or second-line therapy in ALL patients. Pharmacological perturbation of GCN2 suppressed leukemia progression more effectively than ASNase alone, indicating that adaptation to nutrient stress itself constitutes a therapeutic liability. Notably, all tested T-ALL and B-ALL models showed sensitivity to GCN2-IN-6, independent of baseline *ASNS* and *ATF4* levels. GCN2-IN-6 has an additional marked advantage as it is a synthetic compound that will not directly trigger the immune system, unlike the bacterial derived ASNases.^41–43^ Although continuous monotherapy of GCN2-IN-6 with sustained blockage of the AAR pathway was sufficient to efficiently inhibit tumor growth, discontinuation of treatment with the inhibitor eventually resulted in relapse and regrowth of PDX models in immunodeficient NSG mice. This observation suggests that continued stress pathway engagement is required for tumor maintenance and that metabolic adaptation remains highly plastic.

Also in solid PDX models, GCN2-IN-6 had profound anti-tumor effects: in TNBC, ovarian cancer and neuroblastoma GCN2-IN-6 inhibited growth. Interestingly, in neuroblastoma full regression of the tumor could be observed. To obtain complete regression in the other tumor types, further exploration of combination therapies is warranted. Several synergisms have already been identified: Piecyk et al.^44^ found synergy between oxaliplatin and GCN2 inhibition in colon cancer, while Gauthier-Coles et al.^45^ reported synergism with proteasomal inhibitors, such as bortezomib, and CDK inhibitors. Another potential strategy is to combine GCN2 inhibition with drugs further disrupting mitochondrial function, such as sorafenib and nitazonaxide, or with NRF-2 or OXPHOS inhibitors.

In TNBC, heterogeneity in response was observed. This heterogeneity provided an opportunity to investigate determinants of sensitivity. Integrated ribosome profiling and RNA sequencing revealed that sensitive TNBC models exist in a state of chronic metabolic adaptation characterized by constitutive activation of amino acid stress signaling, NRF2-dependent oxidative stress responses and extensive mitochondrial metabolic remodeling. Surprisingly, these tumors exhibited reduced mitochondrial abundance despite enrichment of oxidative phosphorylation pathways and depletion of mitochondrial dysfunction signatures. Rather than indicating mitochondrial impairment, these findings suggest that sensitive tumors maintain substantial oxidative capacity despite a reduced mitochondrial pool.

This phenotype is consistent with an emerging functional connection between ISR activation and mitochondrial adaptation. Vendramin et al.^46^ demonstrated in melanoma that ISR activation suppresses cytosolic protein synthesis while promoting mitochondrial translation, thereby establishing a dependency on mitochondrial protein synthesis for survival under stress. Their findings provide a mechanistic precedent for mitochondrial remodeling as an integral component of the stress-adapted cancer cell state. In the context of our findings, we propose that tumors experiencing chronic metabolic stress coordinately engage amino acid stress signaling, redox defenses, and mitochondrial metabolism to preserve metabolic homeostasis. Such extensive adaptation may simultaneously constrain metabolic flexibility, rendering these tumors particularly vulnerable to additional perturbation of the stress-response network.

In cancer cells, aberrant NRF2 is frequently observed.^47^ Its upregulation helps protect the cancer cells from cellular stress, enhancing their survival potential. Additionally, high NRF2 expression in cancer cells has been linked to resistance to chemotherapy and radiation therapy.^48,49^ High NRF2 expression has also been linked to stabilization of the hybrid E/M state (epithelial/mesenchymal), which can create the plasticity necessary for chemo-evasion.^50^ GCN2 inhibition could thus potentially be a therapy alternative for *NFE2L2*-overexpressing tumors who fail to react well to traditional therapies.

From a translational perspective, our data suggest that constitutive activation of the AAR pathway, as well as mitochondrial function could serve as biomarkers of response. Expression of *ATF4*, *ASNS* and *NFE2L2*, together with reduced mitochondrial DNA content, consistently associated with therapeutic sensitivity in TNBC models. Although these observations require validation in larger cohorts, they provide an initial framework for identifying tumors that are most likely to benefit from therapies targeting stress adaptation.

A major finding of our study is the clear dissociation between genetic suppression of GCN2 and pharmacological treatment with GCN2-IN-6. Genetic knockdown of GCN2 failed to phenocopy the effects of the compound, whereas GCN2-IN-6 induced reproducible transcriptional changes associated with suppression of epithelial-to-mesenchymal transition, extracellular matrix remodeling and stemness-related programs. Moreover, GCN2-IN-6 unexpectedly promoted activation of canonical ISR signaling *in vivo*. These findings are consistent with recent reports demonstrating paradoxical activation of ATP-competitive GCN2 “inhibitors” and indicate that the biological consequences of pharmacological perturbation of this pathway are substantially more complex than originally appreciated. Rather than invalidating the therapeutic observations, these findings emphasize an important conceptual point: off-balancing amino acid stress signaling can expose a profound metabolic vulnerability irrespective of whether the initiating event is canonical GCN2 inhibition. Our data therefore argue that dependency on stress adaptation, rather than dependency on GCN2 itself, represents the critical therapeutic principle.

Several limitations should be acknowledged. The precise molecular target responsible for the anti-tumor effects of GCN2-IN-6 remains incompletely defined, and the contribution of off-target activities and/or secondary cellular responses requires further investigation. Additionally, the studies were performed in immunodeficient models, precluding assessment of how perturbation of amino acid stress signaling affects anti-tumor immunity. Finally, the biomarker signatures identified here require validation across larger and more diverse model systems.

In conclusion, our study identifies chronic metabolic stress adaptation as an actionable vulnerability across multiple aggressive cancer types. We propose that tumors existing in a state of constitutive amino acid stress become dependent on adaptive signaling programs to maintain metabolic homeostasis and are therefore selectively susceptible to pharmacological perturbation of the AAR/ISR network. These findings establish stress adaptation as a therapeutically exploitable state and provide a framework for the development of biomarker-guided strategies targeting metabolic dependencies in cancer.

## Acknowledgments

We thank PDXGhent, OHMX.bio and TRACE PDX platform at KU Leuven for their technical support.

## Author Contributions

Conceptualization and Design; E.P., M.V.T., P.V.V., S.G. Methodology; E.P., I.F., M.V.T., G.M., G.B., S.G., E.L., M.F.B., Data acquisition; E.P., S.T., E.V.A., L.V., K.H., L.M., A.R., K.V.C., F.M., R.B., Analysis; E.P., S.T., I.F., K.T.C., M.V.T., J.D.C., K.V.D.V., PDX model generation; E.P., S.T., K.D., E.L., M.F.B., N.L., Writing Original Draft; E.P. and S.G. Draft Curation: ; E.P., I.F., S.G. Funding Acquisition; E.P., S.G. Supervision; P.N., H.D.R., K.V.D.V., T.L., G.B., P.V.V., S.G.

All authors have read and agreed to the published version of the manuscript.

## Resource availability

Requests for further information and resources should be directed to and will be fulfilled by the lead contact, Steven Goossens. Any additional information required to re-analyze the data reported in this paper is available from the lead contact. There are restrictions to the availability of the used PDX models in this study, especially if there is potential for commercial application. Depending on the PDX model, a materials transfer agreement and/or payment to compensate for the cost for establishment/expansion of the model may be requested.

## Declaration of Interest

G.M. is a co-founder of OHMX.bio, which provides Ribo-seq analysis as a commercial service. Other authors declare no conflicts of interest.

## Funding

This work has been supported by Kom Op Tegen Kanker (KOTK) (project grants to S.G.), the Cancer Research Institute Ghent (CRIG), vzw Kinderkankerfonds (grant to TL, PVV, SG), Ghent University and is part of a Flanders interuniversity consortium IBOF grant (PN, GM and SG). E.P. is a predoctoral fellow with the Research Foundation Flanders (FWO). PN is supported by the Research Foundation Flanders (FWO, G0F4721N, and G0A8B24N). The respective funding sources were not involved in the writing or preparation of this manuscript.

## Declaration of generative AI and AI-assisted technologies in the writing process

During the preparation of this work, the author(s) used PChat GPT-3.5 in order to rephrase certain sentences. After using this tool or service, the author(s) reviewed and edited the content as needed and take(s) full responsibility for the content of the publication.

## Supplementary materials

**FIGURE S1.**
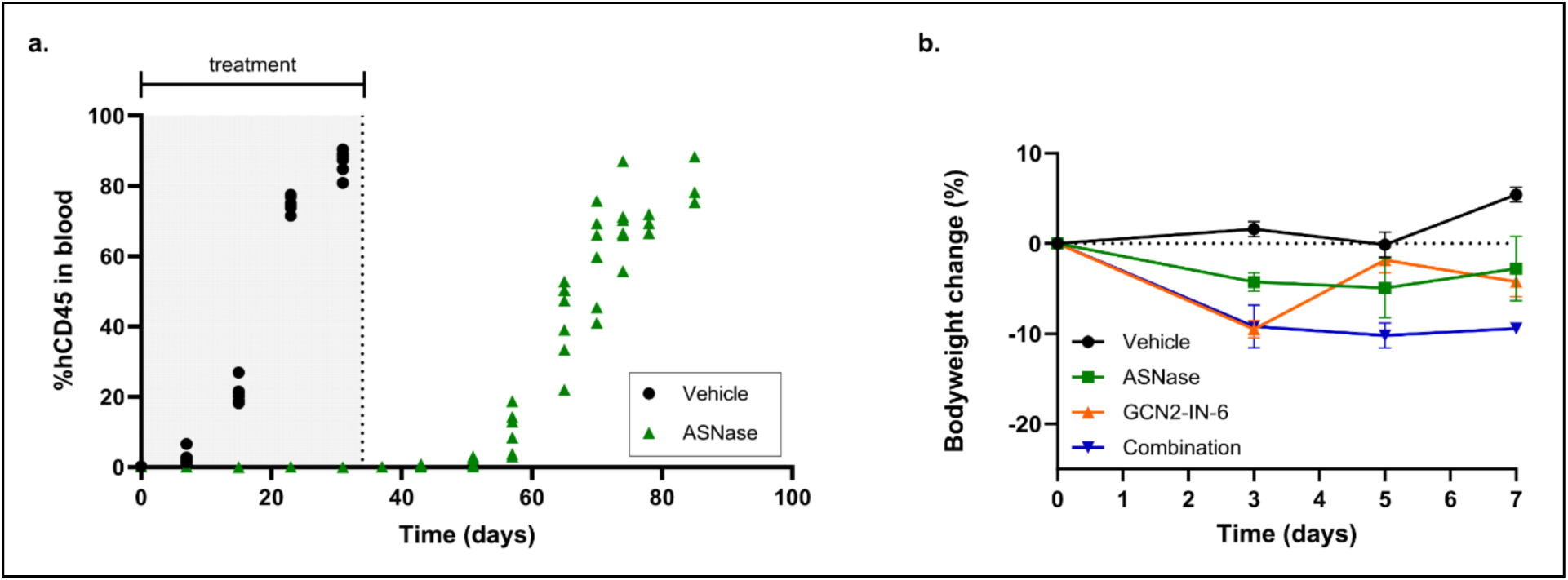
a. Leukemic burden in T-ALL PDX1 mice, measured as the percentage of human CD45-positive (hCD45⁺) cells in peripheral blood during treatment with ASNase (25 IU/mouse every 8 days) for 5 weeks. Treatment was discontinued following sacrifice of the vehicle-treated mice, and the remaining mice were monitored until reaching the experimental endpoint. Each dot represents an individual mouse. b. Bodyweight of mice shown in Figure 1e, expressed as the percentage of baseline bodyweight. Dots represent group means; error bars indicate SEM.

**FIGURE S2.**
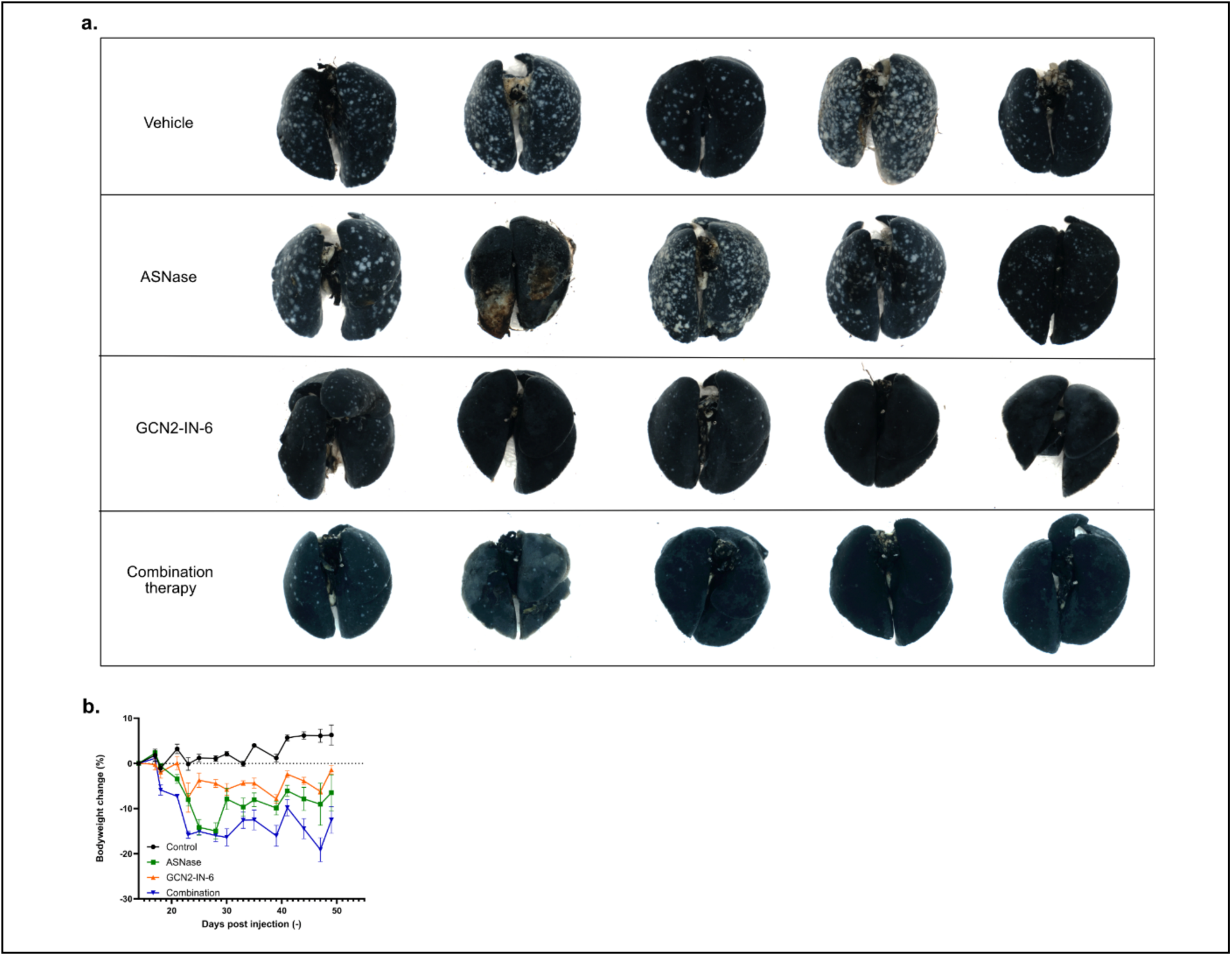
a. Lungs collected from mice shown in Figure 2c-d after intratracheal injection with Indian ink and fixation in Fekete’s solution. Metastatic nodules appear as white lesions against the black-stained lung tissue and were quantified manually. Metastatic burden could not be quantified for one ASNase-treated mouse because of a technical failure during the India ink inflation procedure. b. Body weight of mice shown in Figure 2C, expressed as the percentage of baseline body weight. Dots represent group means; error bars indicate SEM.

**FIGURE S3.**
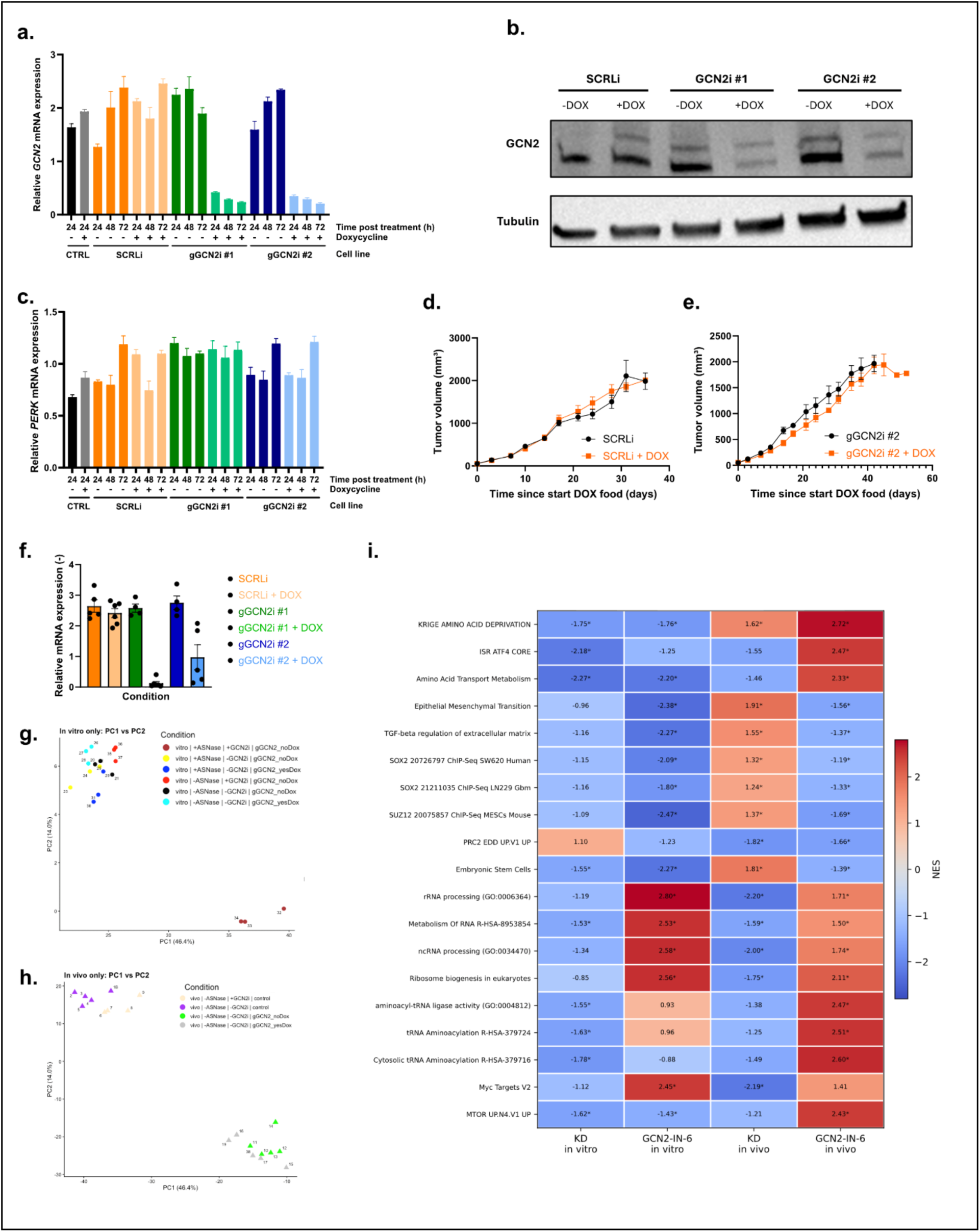
a. Relative *EIF2AK4 (GCN2)* mRNA expression in control (CTRL), scrambled control (gSCRLi), and two independent CRISPRi GCN2 knockdown cell lines (gGCN2i #1 and gGCN2i #2) cultured in the absence or presence of doxycycline (1 μg/mL) for 24, 48, or 72 h. Expression levels were determined by RT-qPCR. Error bars indicate SEM. b. Representative western blot showing GCN2 protein expression in MDAMB231 gSCRLi, gGCN2i #1, and gGCN2i #2 cells cultured with or without doxycycline (1 µg/mL, 48 h). Tubulin serves as the loading control. c. Relative *EIF2AK3 (PERK)* mRNA expression in control (CTRL), scrambled control (gSCRLi), and two independent CRISPRi PERK knockdown cell lines (gGCN2i #1 and gGCN2i #2) cultured in the absence or presence of doxycycline (1 μg/mL) for 24, 48, or 72 h. Expression levels were determined by RT-qPCR. Error bars indicate SEM. d. Tumor volume of orthotopic MDAMB231 gSCRLi xenografts with or without doxycycline induction. Doxycycline was administered continuously in the diet (200 ppm). Dots represent group means (n = 5 mice per group); error bars indicate SEM. e. Tumor volume of orthotopic MDAMB231 gGCN2i #2 xenografts with or without doxycycline-induced GCN2 knockdown. Doxycycline was administered continuously in the diet (200 ppm). Dots represent group means (n = 5 mice per group); error bars indicate SEM. f. Relative *EIF2AK4 (GCN2)* mRNA expression in tumors from the indicated experimental groups at endpoint. Each dot represents an individual tumor; bars indicate mean ± SEM. g. Principal component analysis (PCA) of bulk RNA sequencing data from *in vitro* samples. Experimental groups indicated in figure. h. Principal component analysis (PCA) of bulk RNA sequencing data from *in vivo* tumor samples. Experimental groups indicated in figure. i. Heatmap of normalized enrichment scores (NES) across four experimental contrasts. Candidate gene sets were identified through exploratory Enrichr analysis of genes significantly up- or downregulated following GCN2-IN-6 treatment and selected to represent the principal biological programs identified. Gene sets were subsequently evaluated by preranked GSEA using signed DESeq2 Wald statistics, with weighted enrichment scores calculated using 10,000 gene-set permutations. Permutation-derived values were adjusted for multiple testing using the Benjamini–Hochberg method. Asterisks indicate FDR <0.05.

**FIGURE S4.**
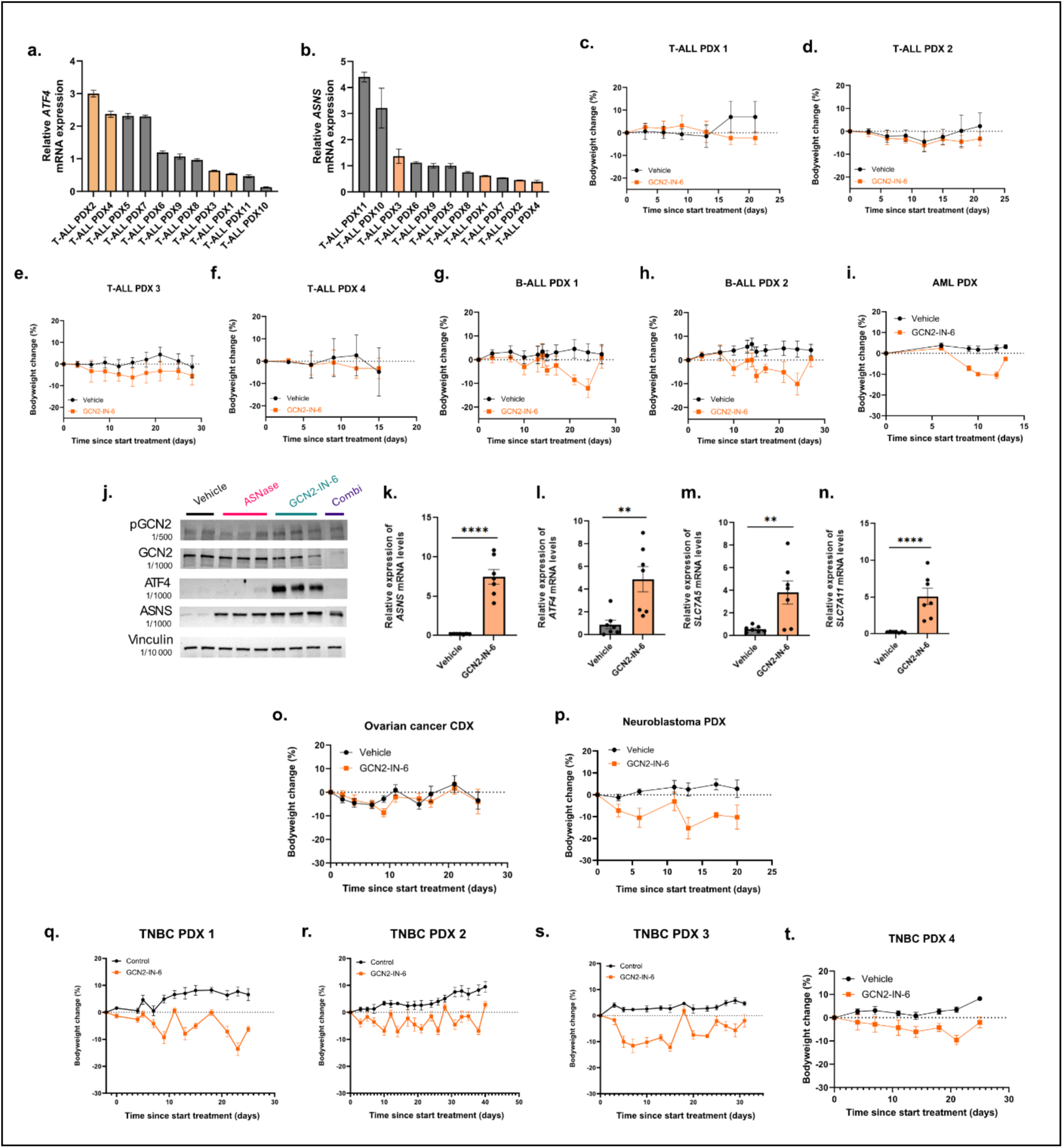
a-b. Relative mRNA expression levels of *ATF4* and *ASNS*, respectively, in different T-ALL PDX models (orange = models selected for *in vivo* treatment with GCN2-IN-6). Error bars denote SEM. c-f. Bodyweight of mice bearing T-ALL PDX models 1–4 during treatment with vehicle or GCN2-IN-6 (3 mg/kg, orally, twice weekly). Bodyweight is expressed as the percentage of change compared to baseline. Error bars indicate SEM. g-h. Bodyweight of mice bearing B-ALL PDX models 1 and 2 during treatment with vehicle or GCN2-IN-6 (3 mg/kg, orally, twice weekly). Bodyweight is expressed as the percentage of change compared to baseline. Error bars indicate SEM. i. Bodyweight of mice bearing the AML PDX during treatment with vehicle or GCN2-IN-6 (3 mg/kg, orally, twice weekly). Bodyweight is expressed as the percentage of change compared to baseline. Error bars indicate SEM. j. Representative western blot analysis of pGCN2, GCN2, ATF4, and ASNS protein expression in tumors harvested from the AML PDX following treatment with vehicle, ASNase, GCN2-IN-6, or the combination. Vinculin serves as the loading control. k-n. Relative mRNA expression of *ASNS*, *ATF4*, *SLC7A5*, and *SLC7A11*, respectively, in tumors derived from AML PDX bearing mice treated with vehicle or GCN2-IN-6 (3 mg/kg, orally, twice weekly). Each dot represents an individual tumor; bars indicate mean ± SEM. Statistical significance was determined using an unpaired two-tailed *t* test. Significance is indicated in the figures. o. Bodyweight of mice bearing the ovarian cancer cell line-derived xenograft (CDX) during treatment with vehicle or GCN2-IN-6 (3 mg/kg, orally, twice weekly). Bodyweight is expressed as the percentage of change compared to baseline. Error bars indicate SEM. p. Bodyweight of mice bearing the neuroblastoma PDX during treatment with vehicle or GCN2-IN-6 (3 mg/kg, orally, twice weekly). Bodyweight is expressed as the percentage of change compared to baseline. Error bars indicate SEM. q-t. Bodyweight of mice bearing TNBC PDX models 1–4 during treatment with vehicle or GCN2-IN-6 (3 mg/kg, orally, twice weekly). Bodyweight is expressed as the percentage of change compared to baseline. Error bars indicate SEM.

**FIGURE S5.**
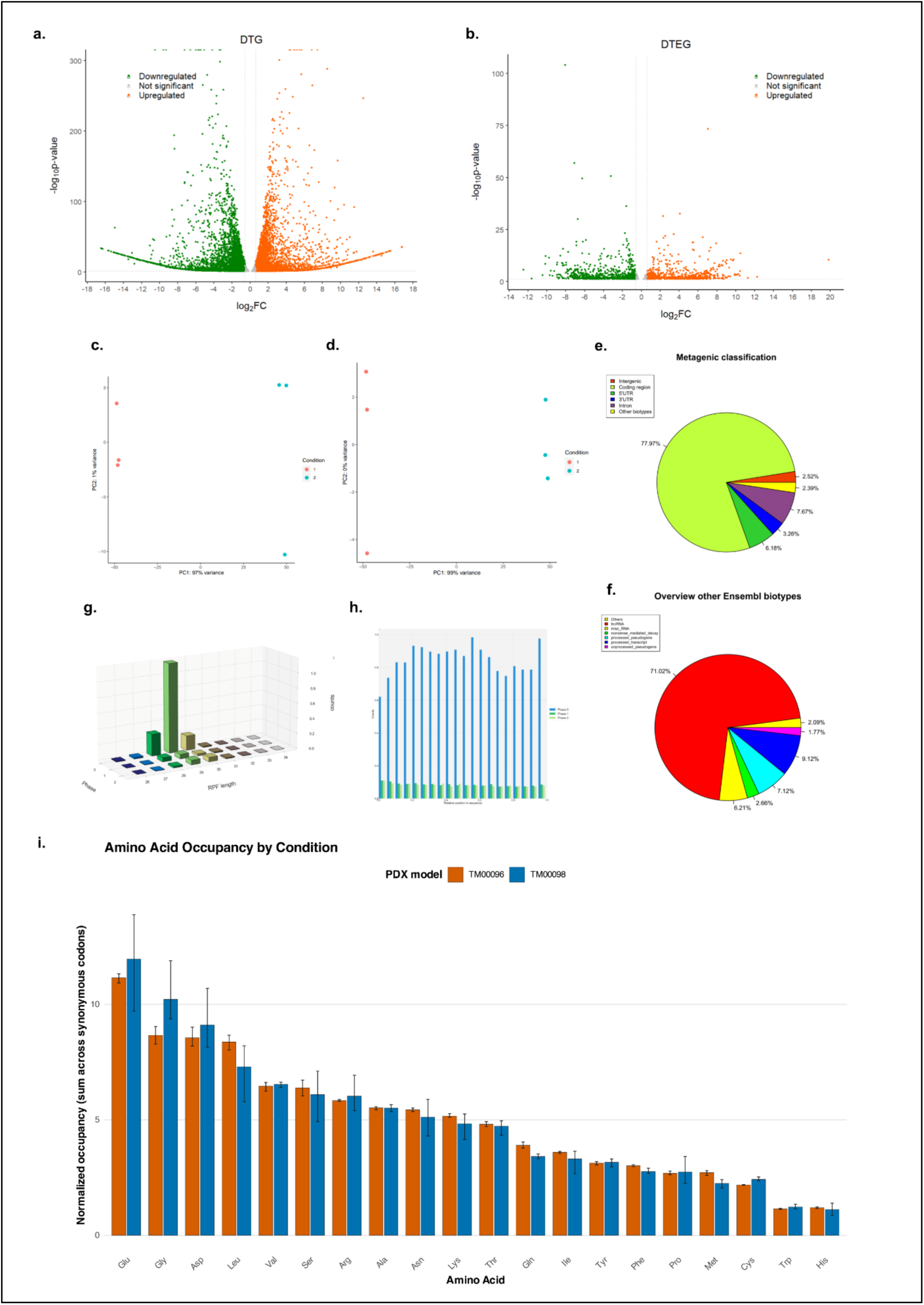
a-b. Volcano plot representing (a) differentially transcribed genes (DTG) and (b) genes with different translation efficiency (DTEG), respectively, from ribosome profiling and RNAseq analysis. Comparison was made between TNBC PDX 1 and TNBC PDX 3, with TNBC PDX 3 used as reference. Significant genes are colored, while non-significant genes are shown in grey. c-d. Principal component analysis (PCA) of (c) RNA-seq and (d) ribosome profiling samples. e. Metagenic distribution of the ribosome protected fragments (RPFs). f. Distribution of reads assigned to non-coding and other Ensembl transcript biotypes (the small pie piece from *e*). g. Length distribution and reading-frame periodicity of ribosome-protected fragments (RPFs). Fragment lengths are stratified according to their inferred reading frame. h. Relative abundancy plot of the RPFs over all coding regions. i. Codon occupancy analysis of all amino acids, comparing TNBC PDX 1 and TNBC PDX 3 models.

**Table S1.** Table representing number of metasiases in lungs. liver and lymph nodes of mice from **Fig 2d**. Counting of metastatic nodules was done manually at time of sac,ifice. Visualisation of lung metastases was done using Indian ink injections.

| **Lung metastases** |  |  |  |
|---|---|---|---|
| **Vehicle** | **PEG-ASNase** | **GCN2-IN-6** | **Combination** |
| 263 | 101 | 0 | 1 |
| 455 | 211 | 1 | 8 |
| 146 | 149 | 30 | 5 |
| 114 | 501 | 93 | 39 |
| 399 | Ink failed | 4 | 3 |
| **Liver metastases** |  |  |  |
| **Vehicle** | **PEG-ASNase** | **GCN2-IN-6** | **Combination** |
| 10 | 1 | 1 | 0 |
| 6 | 0 | 1 | 0 |
| 12 | 1 | 0 | 0 |
| 17 | 0 | 1 | 0 |
| 3 | 1 | 0 | 1 |
| **Lymph node metastases** |  |  |  |
| **Vehicle** | **PEG-ASNase** | **GCN2-IN-6** | **Combination** |
| 1 | 1 | 0 | 0 |
| 1 | 0 | 0 | 0 |
| 0 | 1 | 0 | 0 |
| 3 | 0 | 0 | 0 |
| 0 | 0 | 0 | 0 |

**Table S2.** Table representing the subtype characterization of all leukemic PDX models used. Models are classified by major subtype (T-ALL, B-ALL), before further subclassification is shown.

| **PDX model** | **Subclassification** |
|---|---|
| T-ALL models |  |
| T-ALL PDX 1 | TAL |
| T-ALL PDX 2 | TLX3 |
| T-ALL PDX 3 | TAL |
| T-ALL PDX 4 | TAL |
| T-ALL PDX 5 | NKX2.1 |
| T-ALL PDX 6 | Unknown |
| T-ALL PDX 7 | TLX3 |
| T-ALL PDX 8 | IMM |
| T-ALL PDX 9 | T-LBL TAL |
| T-ALL PDX 10 | Unknown |
| T-ALL PDX 11 | T-LBL NKX2.5 |
| B-ALL models |  |
| B-ALL PDX 1 | none of the major defining genetic lesions (BCR-ABL1,ETV6-RUNX1,high hyperdiploidy, low hypodiploidy, near haploidy, iAMP21, KMT2A-rearrangement, TCF3-PBX1,TCF3-HLF,ABL-class fusions) |
| B-ALL PDX 2 | none of the major defining genetic lesions (BCR-ABL1,ETV6-RUNX1,high hyperdiploidy, low hypodiploidy, near haploidy, iAMP21, KMT2A-rearrangement, TCF3-PBX1,TCF3-HLF,ABL-class fusions) |

**Table S3.**
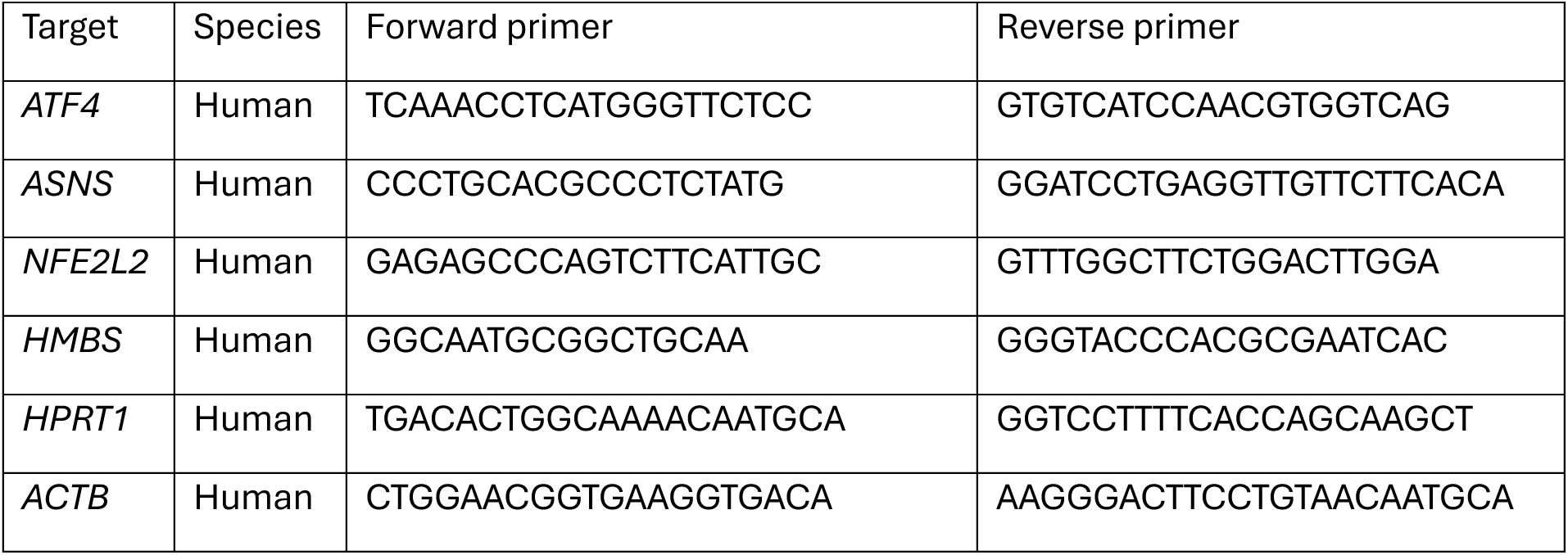
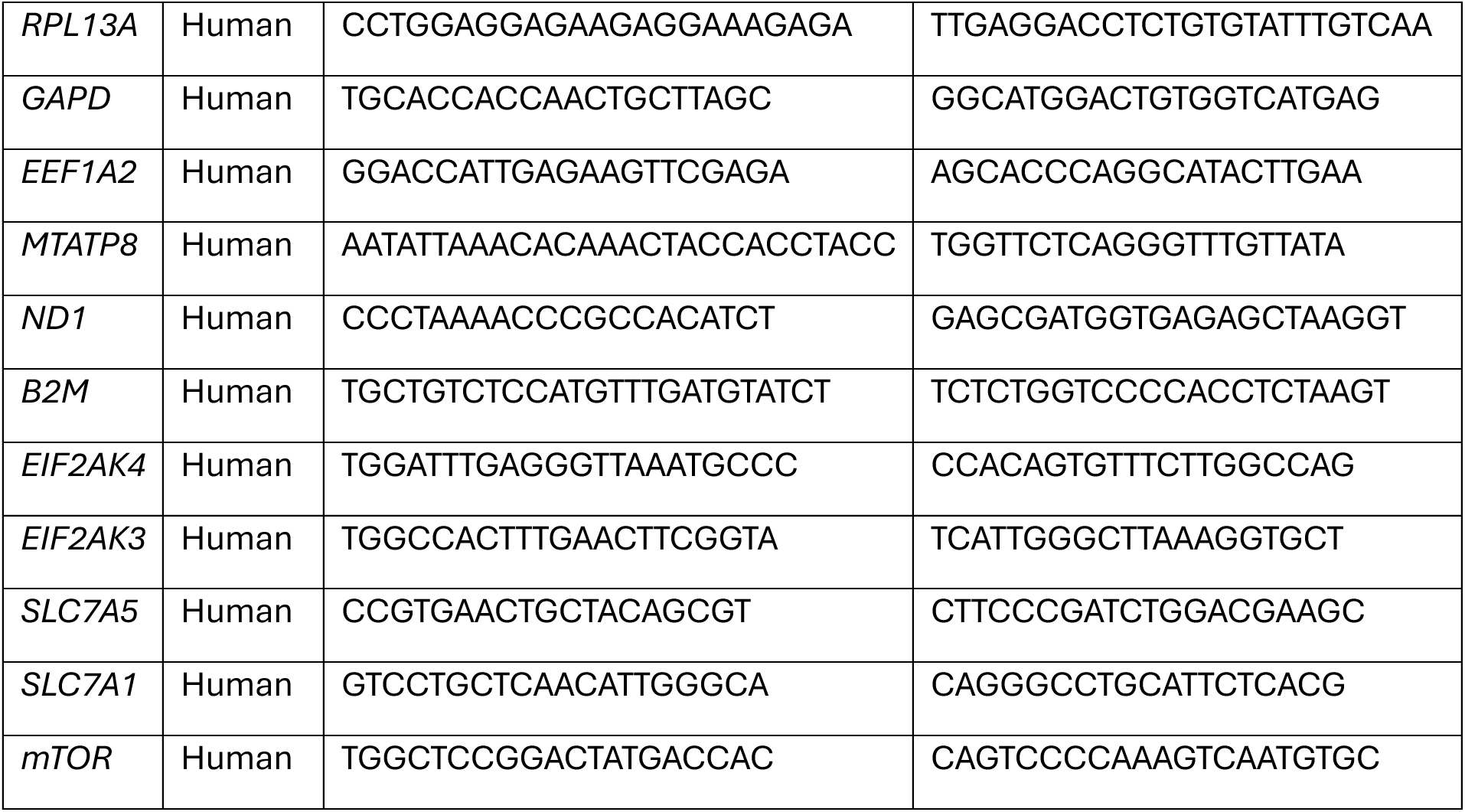

**Table S4.** Filtering and mapping statistics for the RIBOseq data (green annotation = optimal, red annotation = problematic)

| sample | type | total | mapped_U | mapped_M | mapped_T | unmapped | map_freq_U | map_freq_M | map_freq_T | map_freq_total |
|---|---|---|---|---|---|---|---|---|---|---|
| OHMX20230139_001_STAR_Y_105.fastq1 | clipper | 50.557.332 |  |  | 48.379.374 | 2.177.958 |  |  | 95,69% |  |
| OHMX20230139_001_STAR_Y_105.fastq1 | rRNA | 48.379.374 |  |  | 4.720.424 | 43.658.950 |  |  | 9,76% |  |
| OHMX20230139_001_STAR_Y_105.fastq1 | snRNA | 43.658.950 |  |  | 2.968.501 | 40.690.449 |  |  | 6,80% |  |
| OHMX20230139_001_STAR_Y_105.fastq1 | tRNA | 40.690.449 |  |  | 1.058.394 | 39.632.055 |  |  | 2,60% |  |
| OHMX20230139_001_STAR_Y_105.fastq1 | genomic | 39.632.055 | 24.845.360 | 0 | 24.845.360 | 14.786.695 | 62,69% | 0,00% | 62,69% | 51,36% |
| OHMX20230139_002_STAR_Y_105.fastq1 | clipper | 67.003.477 |  |  | 57.536.415 | 9.467.062 |  |  | 85,87% |  |
| OHMX20230139_002_STAR_Y_105.fastq1 | rRNA | 57.536.415 |  |  | 4.109.376 | 53.427.039 |  |  | 7,14% |  |
| OHMX20230139_002_STAR_Y_105.fastq1 | snRNA | 53.427.039 |  |  | 1.799.287 | 51.627.752 |  |  | 3,37% |  |
| OHMX20230139_002_STAR_Y_105.fastq1 | tRNA | 51.627.752 |  |  | 600.897 | 51.026.855 |  |  | 1,16% |  |
| OHMX20230139_002_STAR_Y_105.fastq1 | genomic | 51.026.855 | 32.885.549 | 0 | 32.885.549 | 18.141.306 | 64,45% | 0,00% | 64,45% | 57,16% |
| OHMX20230139_003_STAR_Y_105.fastq1 | clipper | 60.570.443 |  |  | 56.241.363 | 4.329.080 |  |  | 92,85% |  |
| OHMX20230139_003_STAR_Y_105.fastq1 | rRNA | 56.241.363 |  |  | 7.425.556 | 48.815.807 |  |  | 13,20% |  |
| OHMX20230139_003_STAR_Y_105.fastq1 | snRNA | 48.815.807 |  |  | 3.169.704 | 45.646.103 |  |  | 6,49% |  |
| OHMX20230139_003_STAR_Y_105.fastq1 | tRNA | 45.646.103 |  |  | 930.893 | 44.715.210 |  |  | 2,04% |  |
| OHMX20230139_003_STAR_Y_105.fastq1 | genomic | 44.715.210 | 24.573.039 | 0 | 24.573.039 | 20.142.171 | 54,95% | 0,00% | 54,95% | 43,69% |
| OHMX20230139_004_STAR_Y_105.fastq1 | clipper | 58.292.543 |  |  | 56.161.699 | 2.130.844 |  |  | 96,34% |  |
| OHMX20230139_004_STAR_Y_105.fastq1 | rRNA | 56.161.699 |  |  | 2.526.202 | 53.635.497 |  |  | 4,50% |  |
| OHMX20230139_004_STAR_Y_105.fastq1 | snRNA | 53.635.497 |  |  | 587.430 | 53.048.067 |  |  | 1,10% |  |
| OHMX20230139_004_STAR_Y_105.fastq1 | tRNA | 53.048.067 |  |  | 1.609.398 | 51.438.669 |  |  | 3,03% |  |
| OHMX20230139_004_STAR_Y_105.fastq1 | genomic | 51.438.669 | 35.152.420 | 0 | 35.152.420 | 16.286.249 | 68,34% | 0,00% | 68,34% | 62,59% |
| OHMX20230139_005_STAR_Y_105.fastq1 | clipper | 59.460.765 |  |  | 58.546.503 | 914.262 |  |  | 98,46% |  |
| OHMX20230139_005_STAR_Y_105.fastq1 | rRNA | 58.546.503 |  |  | 3.286.619 | 55.259.884 |  |  | 5,61% |  |
| OHMX20230139_005_STAR_Y_105.fastq1 | snRNA | 55.259.884 |  |  | 646.909 | 54.612.975 |  |  | 1,17% |  |
| OHMX20230139_005_STAR_Y_105.fastq1 | tRNA | 54.612.975 |  |  | 577.693 | 54.035.282 |  |  | 1,06% |  |
| OHMX20230139_005_STAR_Y_105.fastq1 | genomic | 54.035.282 | 37.786.471 | 0 | 37.786.471 | 16.248.811 | 69,93% | 0,00% | 69,93% | 64,54% |
| OHMX20230139_006_STAR_Y_105.fastq1 | clipper | 53.447.097 |  |  | 49.412.481 | 4.034.616 |  |  | 92,45% |  |
| OHMX20230139_006_STAR_Y_105.fastq1 | rRNA | 49.412.481 |  |  | 2.440.724 | 46.971.757 |  |  | 4,94% |  |
| OHMX20230139_006_STAR_Y_105.fastq1 | snRNA | 46.971.757 |  |  | 422.550 | 46.549.207 |  |  | 0,90% |  |
| OHMX20230139_006_STAR_Y_105.fastq1 | tRNA | 46.549.207 |  |  | 1.771.168 | 44.778.039 |  |  | 3,80% |  |
| OHMX20230139_006_STAR_Y_105.fastq1 | genomic | 44.778.039 | 27.992.370 | 0 | 27.992.370 | 16.785.669 | 62,51% | 0,00% | 62,51% | 56,65% |
| OHMX20230139_007_STAR_Y_105.fastq1 | clipper | 51.909.964 |  |  | 50.032.386 | 1.877.578 |  |  | 96,38% |  |
| OHMX20230139_007_STAR_Y_105.fastq1 | rRNA | 50.032.386 |  |  | 3.628.561 | 46.403.825 |  |  | 7,25% |  |
| OHMX20230139_007_STAR_Y_105.fastq1 | snRNA | 46.403.825 |  |  | 730.122 | 45.673.703 |  |  | 1,57% |  |
| OHMX20230139_007_STAR_Y_105.fastq1 | tRNA | 45.673.703 |  |  | 320.710 | 45.352.993 |  |  | 0,70% |  |
| OHMX20230139_007_STAR_Y_105.fastq1 | genomic | 45.352.993 | 31.726.700 | 0 | 31.726.700 | 13.626.293 | 69,96% | 0,00% | 69,96% | 63,41% |
| OHMX20230139_008_STAR_Y_105.fastq1 | clipper | 54.245.554 |  |  | 51.149.894 | 3.095.660 |  |  | 94,29% |  |
| OHMX20230139_008_STAR_Y_105.fastq1 | rRNA | 51.149.894 |  |  | 3.383.725 | 47.766.169 |  |  | 6,62% |  |
| OHMX20230139_008_STAR_Y_105.fastq1 | snRNA | 47.766.169 |  |  | 931.795 | 46.834.374 |  |  | 1,95% |  |
| OHMX20230139_008_STAR_Y_105.fastq1 | tRNA | 46.834.374 |  |  | 258.257 | 46.576.117 |  |  | 0,55% |  |
| OHMX20230139_008_STAR_Y_105.fastq1 | genomic | 46.576.117 | 30.338.712 | 0 | 30.338.712 | 16.237.405 | 65,14% | 0,00% | 65,14% | 59,31% |
| OHMX20230139_009_STAR_Y_105.fastq1 | clipper | 56.581.341 |  |  | 53.609.323 | 2.972.018 |  |  | 94,75% |  |
| OHMX20230139_009_STAR_Y_105.fastq1 | rRNA | 53.609.323 |  |  | 5.359.708 | 48.249.615 |  |  | 10,00% |  |
| OHMX20230139_009_STAR_Y_105.fastq1 | snRNA | 48.249.615 |  |  | 836.290 | 47.413.325 |  |  | 1,73% |  |
| OHMX20230139_009_STAR_Y_105.fastq1 | tRNA | 47.413.325 |  |  | 332.237 | 47.081.088 |  |  | 0,70% |  |
| OHMX20230139_009_STAR_Y_105.fastq1 | genomic | 47.081.088 | 30.324.631 | 0 | 30.324.631 | 16.756.457 | 64,41% | 0,00% | 64,41% | 56,57% |

## Materials and methods

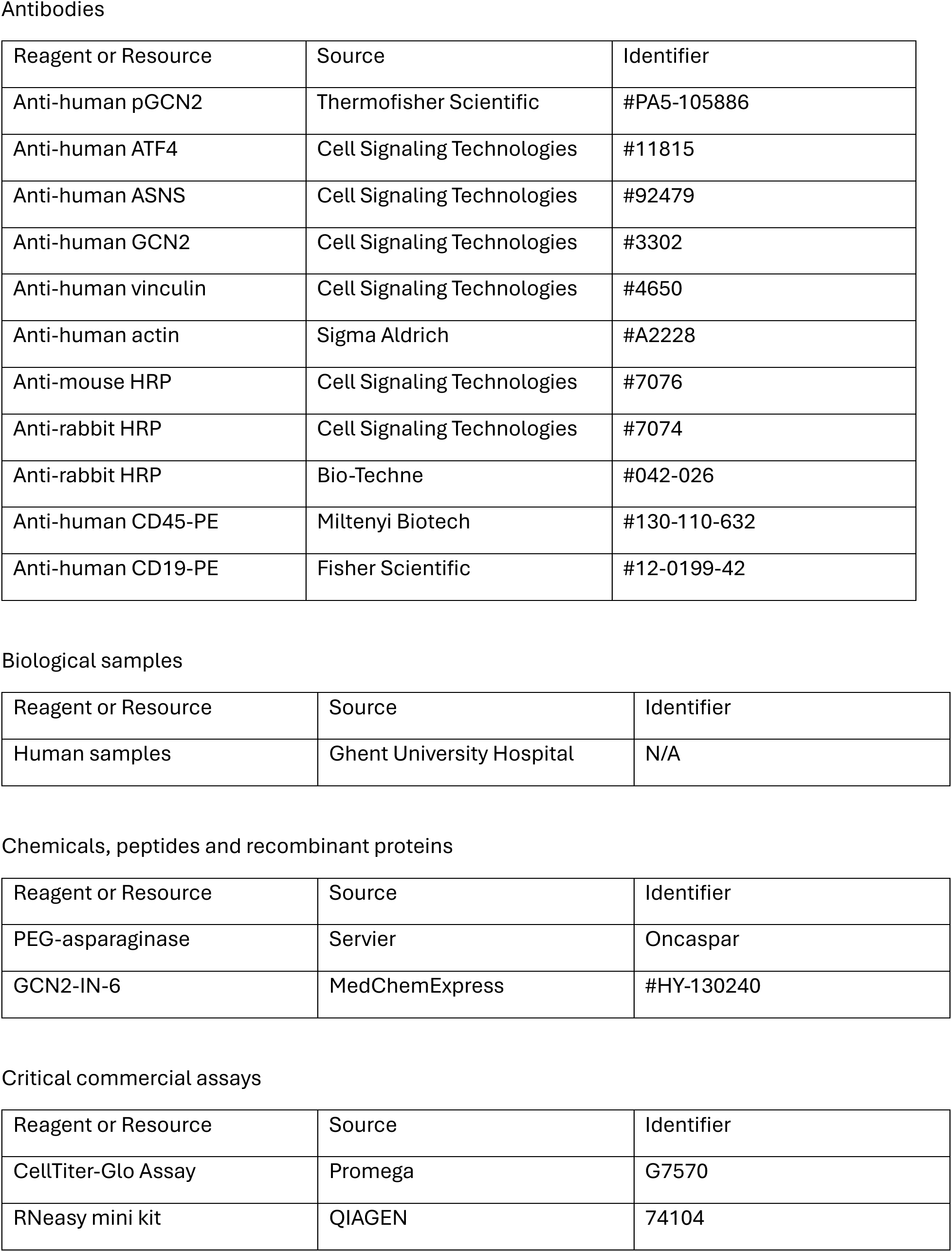
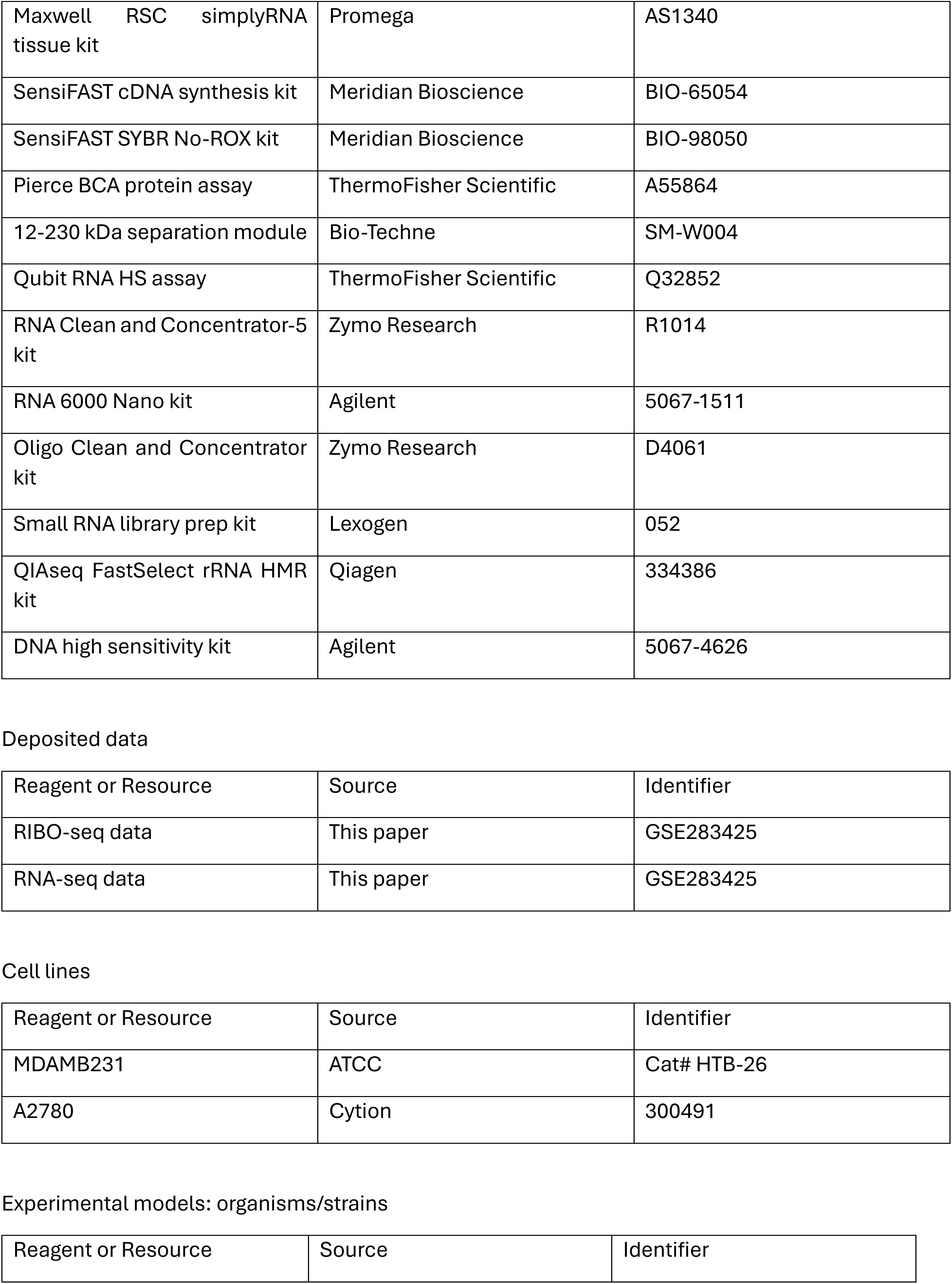
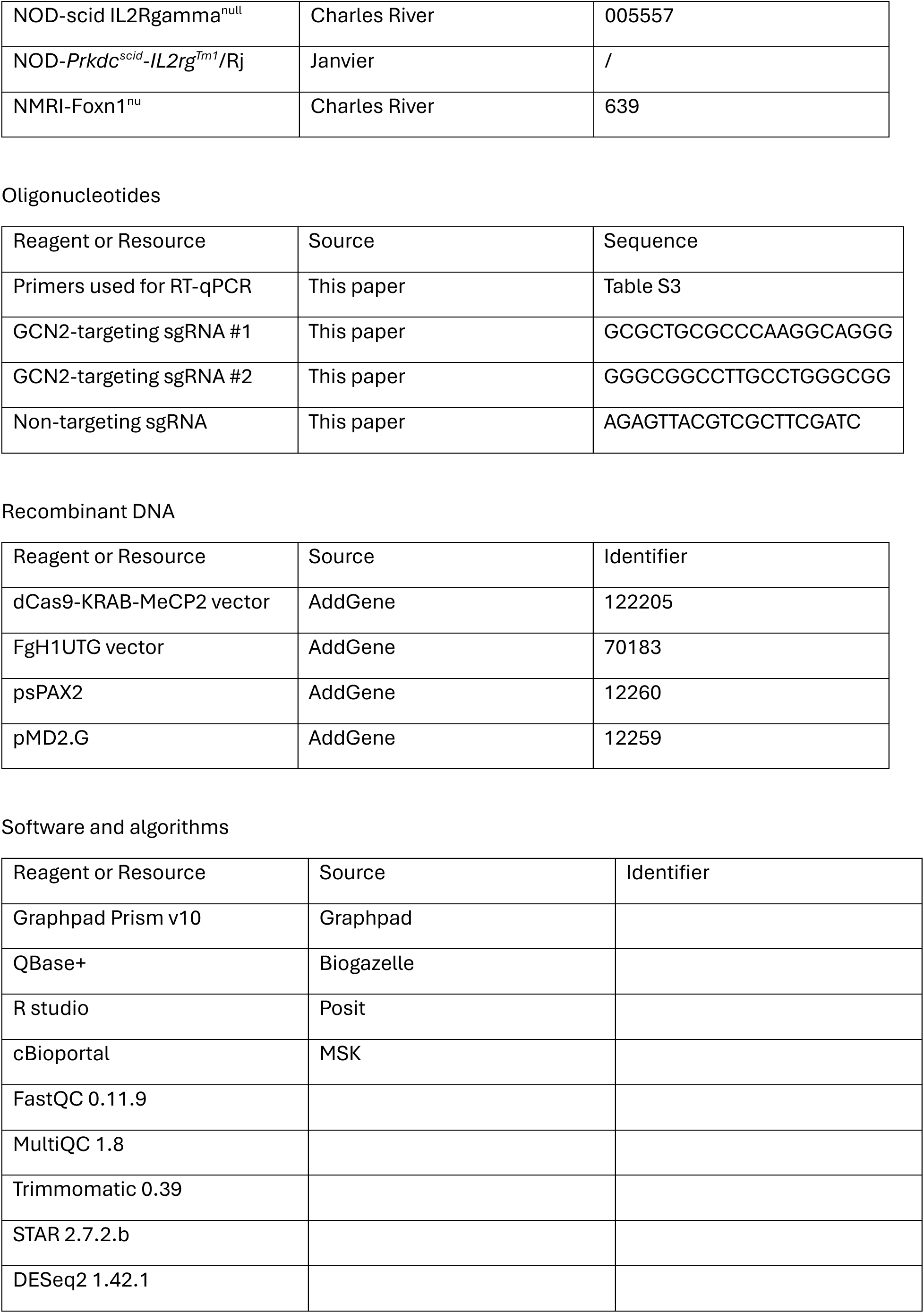

## Resource availability

### Lead contact

Further information and requests for resources and reagents should be directed to and will be fulfilled by the lead contact, Steven Goossens.

### Materials availability

This study did not generate new unique reagents.

### Data and code availability

- Matched RIBOseq and RNAseq data analyzed in this study have been deposited at GEO (GSE283425). They are publicly available at the time of publication and are listed in the key resources table. Freely available software and algorithms used for analysis are also listed in the key resources table.

- RNAseq data has been deposited at GEO (GSE343685).

- This paper does not report original code.

- Any additional information required to reanalyze the data reported in this work paper is available from the lead contact upon reasonable request.

## Experimental models

### Cell culture

MDAMB231 breast cancer cells were obtained from ATCC (HTB-26) and were cultured in high-glucose DMEM (Gibco, #11965-092), supplemented with 10% fetal calf serum, L-glutamine, sodium pyruvate and non-essential amino acids. Human A2780 ovarian cancer cells were obtained from Cytion (#300491) and cultured in RPMI 1640 (Gibco, #52400-025), supplemented with 10% fetal calf serum and L-glutamine. All cell lines were incubated at 37°C in a 5% CO_2_ humidified environment.

### Patient-derived leukemia cells

Patient-derived leukemia cells (PDCs) were obtained from the spleens of mice bearing patient-derived leukemia xenografts. Mice were euthanized when peripheral blood leukemic burden exceeded 80% human CD45-positive (hCD45⁺) cells. Spleens were harvested and mechanically dissociated through a 70-μm cell strainer. Red blood cells were removed using ACK lysis buffer (Gibco, #A1049201), after which cells were resuspended in RPMI 1640 medium (Gibco, #52400-025) supplemented with 10% fetal calf serum and 1% penicillin-streptomycin. Cells were maintained at 37°C in a humidified atmosphere containing 5% CO₂.

### PDX models

Breast cancer PDX models TM00098 (BC PDX 1) and TM00096 (BC PDX 3) were commercially obtained from the Jackson Laboratory and passaged in-house. Breast cancer PDX BRC0004 (BC PDX 2) was developed by the TRACE PDX Repository (KULeuven). Breast cancer TNBC PDX 4, all leukemic PDX models and the AML PDX were created from patient material at Ghent University Hospital under approved ECDs. The neuroblastoma NEC005 PDX model was obtained through collaboration with the Experimental Pharmacology and Oncology (EPO) lab (Berlin, Germany) and expanded at Ghent University Hospital through subcutaneous injection into NSG mice. All experiments with BRC0004 were performed by TRACE in the KULeuven animal facility. All other experiments were performed at MRB2 animal facility at Ghent University Hospital.

### Animals

Animal experiments were performed at Ghent University and KU Leuven and were approved by the respective institutional animal ethics committees. All procedures were conducted in accordance with applicable institutional, national, and European regulations governing animal research.

Mice were housed in ventilated cages in specific pathogen-free, environmentally controlled facilities at 21°C–25°C under a 12h light/dark cycle, with food and water provided ad libitum. Animals were group-housed with up to five mice per cage. Bedding, nesting material, and shelters were provided as environmental enrichment.

NOD/SCID/gamma (NSG) mice were obtained from Charles River (#005557). NOD-Prkdc^scid^-IL2rg^Tm^^1^/Rj (NXG) mice were obtained from Janvier.

Animals were randomly allocated to treatment groups after establishment of disease or measurable tumor growth, as described for the individual experiments.

### Reagents C compounds

Oncaspar (PEG-ASNase) was obtained from a commercial supplier (Servier). GCN2-IN-6 was obtained from a commercial supplier (Medchem, #HY-130240).

Oncaspar was dissolved in physiological water for both *in vitro* and *in vivo* experiments.

For *in vitro* experiments, GCN2-IN-6 was dissolved in dimethyl sulfoxide (DMSO) to prepare a stock solution and subsequently diluted in culture medium to the indicated working concentrations.

For *in vivo* administration, GCN2-IN-6 was formulated in a vehicle consisting of 10% DMSO, 40% PEG300, 5% Tween-80, and 45% saline and administered orally at the indicated dose and schedule.

### Method details

#### Cell viability assays

Cell viability was assessed using CellTiter-Glo Luminescent viability assay (Promega, #G7570). Cells were seeded in white 96-well plates at a density of 400 cells/well in the appropriate culture medium and allowed to adhere for 24 hours at 37°C in a humidified atmosphere containing 5% CO_2_. Cells were subsequently treated with ASNase (0 – 10 IU/mL, 2x dilution) and/or GCN2-IN-6 (0 – 2000 nM, 2x dilution) for 72 hours. Following treatment, CellTiter-Glo reagent was prepared according to the manufacturer’s instructions and added to each well at a ratio of 1:2 reagens:medium. Plates were mixed for 2 min at 200 rpm, followed by incubation at room temperature for 10 minutes, before luminescence was measured. Read-out of luminescence was performed using a GloMax Explorer luminometer (Promega, #GM3500).

Cell viability was expressed relative to vehicle-treated cells, which were set to 100%. Dose-response curves and IC₅₀ values were determined by nonlinear regression using GraphPad Prism v10.

#### Ex vivo PEG-ASNase sensitivity of leukemia cells

Leukemia cells were isolated from the spleens of T-ALL PDX 1-bearing mice five days after administration of a single 100 IU dose of PEG-ASNase or vehicle, as described below. Isolated cells were treated *ex vivo* with a two-fold serial dilution of PEG-ASNase up to a maximum concentration of 2,000 mIU/mL for 72 h. Cell viability was subsequently assessed using CellTiter-Glo as described above.

#### Ǫuantitative real-time PCR

Total cellular RNA was extracted using the RNeasy mini kit (ǪIAGEN, #74104) or the Maxwell RSC simplyRNA tissue kit (Promega, #AS1340) according to the manufacturer’s protocols. The cDNA was synthesized through reverse transcription of RNA using the SensiFAST cDNA synthesis kit (Meridian Bioscience, #BIO-65054). Ǫuantitative real-time PCR was performed using the SensiFAST SYBR No-ROX kit (Meridian Bioscience, #BIO-98050) on a LightCycler 480 II system. All primers that were used can be found in *Table S3*.

#### Mitochondrial DNA quantification

Relative mitochondrial DNA content was determined by quantitative PCR. Snap-frozen primary tumor tissue was homogenized using a TissueLyser for 5 min at 20 Hz. DNA was subsequently isolated using the DNeasy Blood C Tissue Kit (ǪIAGEN, #69504) according to the manufacturer’s instructions.

Mitochondrial DNA abundance was determined using primers targeting *MTATP8* and *ND1*. Genomic DNA abundance was determined using primers targeting *B2M* and *GAPDH*. The mean signal of *MTATP8* and *ND1* was normalized to the mean signal of *B2M* and *GAPDH*, and relative mitochondrial DNA content was expressed as the mtDNA/gDNA ratio. Primer sequences are provided in *Table S3*.

### Western blot

Cells and tumor tissue were lysed in 1% NP-40 buffer (50 mM Tris (pH 7.4), 150 mM NaCl, 1% NP-40), supplemented with Halt protease C phosphatase inhibitor cocktail (ThermoFisher Scientific, #78446) and stored at -80°C prior to processing for protein extracts. Protein concentrations were determined using the Pierce BCA Protein Assay (Thermo Fisher Scientific, #A55864) according to the manufacturer’s instructions.

For conventional western blotting, protein samples were denatured at 95°C in Laemmli sample buffer supplemented with β-mercaptoethanol and separated on 10% polyacrylamide gels in Tris/glycine/SDS running buffer (Bio-Rad). Proteins were transferred to nitrocellulose membranes, which were blocked for 1 h at room temperature in 5% non-fat milk or bovine serum albumin in TBS-T (Tris-buffered saline containing 0.1% Tween-20).

Membranes were incubated overnight at 4°C with the following primary antibodies: anti-phospho-GCN2 (Thermo Fisher Scientific, #PA5-105886; 1:500), anti-ATF4 (Cell Signaling Technology, #11815; 1:1000), anti-ASNS (Cell Signaling Technology, #92479; 1:1000), anti-vinculin (Cell Signaling Technology, #4650; 1:10000), and anti-GCN2 (Cell Signaling Technologies, #3302; 1:1000).

Following washing in TBS-T, membranes were incubated with the appropriate HRP-conjugated secondary antibodies for 45 min at room temperature. Chemiluminescent signal was developed using SuperSignal West Dura (Thermo Fisher Scientific, #34075) or SuperSignal West Femto (Thermo Fisher Scientific, #34095), depending on signal intensity, and visualized using an iBright 750 imaging system (Thermo Fisher Scientific)

### Automated western blotting (WES)

Cells and tumor tissue were lysed in 1% NP-40 buffer (50 mM Tris (pH 7.4), 150 mM NaCl, 1% NP-40), supplemented with Halt protease C phosphatase inhibitor cocktail (ThermoFisher Scientific, #78446) and stored at -80°C prior to processing for protein extracts. Protein concentrations were determined using the Pierce BCA Protein Assay (Thermo Fisher Scientific, #A55864) according to the manufacturer’s instructions.

Automated capillary western blotting was performed using the Wes system (Bio-Techne) and the 12–230 kDa separation module (#SM-W004) according to the manufacturer’s instructions.

Primary antibodies were anti-ASNS (Cell Signaling Technology, #92479; 1:100) and anti-actin (Sigma-Aldrich, #A2228; 1:100). HRP-conjugated anti-rabbit (Bio-Techne, #042-026; used undiluted) and anti-mouse (Cell Signaling Technology, #7076; 1:100) antibodies were used as secondary antibodies. Chemiluminescent signals were detected using the Wes imaging system (Bio-Techne, #004-600).

### CRISPR interference

MDAMB231 cells were engineered to stably express a nuclease-deficient Cas9 fused to the Krüppel-associated box (dCas9-KRAB-MeCP2) transcriptional repressor to enable CRISPR interference (CRISPRi)-mediated gene silencing. Cells were first transduced with a lentiviral vector encoding dCas9-KRAB-MeCP2 (AddGene, #122205) and selected using 8 µg/mL blasticidin to establish a stable CRISPRi parental cell population.

Single-guide RNAs (sgRNAs) targeting EIF2AK4 (GCN2) were designed using CRISPick and cloned into a lentiviral sgRNA expression vector (FgH1tUTG; AddGene, #70183). A non-targeting scrambled sgRNA was used as a negative control. For the sequences, see reagent table.

Lentiviral particles were produced in HEK293T cells by co-transfection of the transfer plasmid with the packaging plasmids psPAX2 and pMD2.G using JetPEI mix according to the manufacturer’s instructions. Viral supernatants were collected 48 and 72 h after transfection, filtered through a 0.45 µm filter, and used to transduce dCas9-KRAB-expressing MDA-MB-231 cells in the presence of 8 µg/mL polybrene. Successfully transduced cells were selected using GFP signal to generate stable polyclonal cell populations.

Two independent GCN2-targeting CRISPRi cell populations (gGCN2i #1 and gGCN2i #2) and a non-targeting control population (gSCRLi) were generated.

For inducible CRISPRi experiments, gene silencing was induced using doxycycline at 1 μg/mL for the indicated duration. Knockdown efficiency was confirmed by RT-qPCR and western blotting before functional experiments.

### *In vivo* experiments

#### General information about *in vivo* experiments

All animal experiments were conducted at Ghent University and KULeuven and approved by the respective animal ethical committees. The studies were conducted in strict accordance with all relevant ethical regulations regarding animal research. Animals were randomly housed in groups, with up to 5 mice per cage. They were housed in ventilated cages in a pathogen-free, environment-controlled room at 21°C-25°C with a 12h light/dark cycle. Water and food was provided *ad libitum*. Cages contained paper chip bedding and a house and nest material were provided as form of enrichment.

#### *In vivo* treatment of T-ALL PDX 1 with PEG-ASNase

T-ALL PDX1 cells (5 × 10⁶ cells/mouse) were injected intravenously into 12 female NSG mice aged 11–14 weeks. Leukemic engraftment was monitored by measuring the percentage of hCD45⁺ cells in peripheral blood. Treatment was initiated once peripheral blood leukemic burden exceeded 1% hCD45⁺ cells.

Mice were randomized to receive PEG-ASNase (25 IU/mouse, intraperitoneally every 8 days; n = 6) or vehicle (n = 6). Mice were sacrificed when %hCD45 reached the threshold of 80%, when bodyweight loss of more than 20% was observed, or in case of general health problems, such as cessation of drinking, eating or nest building. Once all control mice had been sacrificed, PEG-ASNase treatment was also stopped and mice were further followed up for survival. Mice were euthanized when peripheral blood leukemic burden reached 80% hCD45⁺ cells, bodyweight loss exceeded 20% relative to baseline, or predefined clinical humane endpoints were reached, including cessation of eating, drinking, or nest-building behavior. When all vehicle-treated mice had reached experimental endpoint, PEG-ASNase treatment was discontinued and remaining animals were monitored for survival.

#### Short-term PEG-ASNase treatment of T-ALL PDX1 prior to *ex vivo* drug-sensitivity assays

T-ALL PDX1 cells (1.5 × 10⁶ cells/mouse) were intravenously injected into six male NSG mice aged 21–24 weeks. Once peripheral blood leukemic burden reached at least 50% hCD45⁺ cells in all animals, mice were randomly allocated to two treatment groups (n = 3/group) and received a single dose of PEG-ASNase (100 IU/mouse) or vehicle.

Five days after treatment, mice were euthanized and spleens were collected. Splenic tissue was mechanically dissociated twice through a 70-μm cell strainer, followed by red blood cell removal using ACK lysis buffer for 5 min at room temperature. Isolated leukemia cells were subsequently used for ex vivo drug-sensitivity assays.

#### *In vivo* combination treatment in T-ALL PDX 1

T-ALL PDX1 cells (5 × 10⁶ cells/mouse) were injected intravenously into 12 female NSG mice aged 11–14 weeks. Mice were allocated to vehicle, PEG-ASNase, GCN2-IN-6, or combination treatment groups (n = 3/group). PEG-ASNase was administered once intraperitoneally at 25 IU/mouse, and GCN2-IN-6 was administered orally at 3 mg/kg every 3.5 days. Mice were euthanized 7 days after initiation of treatment. Spleens were collected and processed to obtain leukemia cells for downstream analyses.

#### *In vivo* compound treatment of other leukemic PDX models

For each leukemia PDX model, 5 × 10⁶ cells were injected intravenously into 12 female NSG mice aged 6–8 weeks. Leukemic engraftment was monitored by flow cytometric quantification of human leukemia cells in peripheral blood.

Treatment was initiated upon evidence of leukemia engraftment, defined as >1% hCD45⁺ cells for T-ALL and AML or >1% hCD19⁺ cells for B-ALL. Mice were randomized into two groups (n = 6/group) receiving either GCN2-IN-6 (3 mg/kg, oral administration every 4 days) or vehicle. When vehicle-treated mice reached the experimental endpoint, GCN2-IN-6 treatment was discontinued and remaining animals were monitored for survival.

Mice were euthanized when peripheral blood leukemic burden reached 80% hCD45⁺ cells for T-ALL or 40% hCD19⁺ cells for B-ALL, when bodyweight loss exceeded 20% relative to baseline, or when predefined clinical humane endpoints were reached.

#### Flowcytometric assessment of leukemia burden

Peripheral blood was collected via a small cut in the tail vein in EDTA-coated tubes at the indicated time points. Red blood cells were lysed using ACK lysis buffer (Gibco, #A1049201) , and remaining cells were stained using antibodies against human CD45 (Miltenyi Biotech, #130-110-632) or human CD19 (#12-0199-42).

Samples were acquired using the BD FACSymphony™ A3 Cell Analyzer and analyzed using *FlowJo*™ Software v1.

#### *In vivo* combination therapy in MDAMB231 CDX model

MDAMB231 cells were harvested and resuspended in PBS. A total of 3 × 10⁶ cells in 50 μL PBS were orthotopically injected into the fourth mammary fat pad of female NSG mice using a 30-G needle under general anesthesia. The surgical incision was closed using simple sutures, and postoperative analgesia was provided by subcutaneous administration of 100 μL Temgesic (buprenorphine; 0.05 mg/kg).

Tumor dimensions were measured using a caliper and tumor volume was calculated as (length x width²)/2. Once all primary tumors were measurable (volume > 62.5 mm³), the mice were randomly divided into 4 treatment groups (n = 5 per group): vehicle, PEG-ASNase, GCN2-IN-6, or the combination. PEG-ASNase was administered intraperitoneally every 8 days at 25 IU/mouse. GCN2-IN-6 was administered orally every 4 days at a concentration of 3 mg/kg.

When the first animal reached a primary tumor volume > 2000 mm³, all mice were euthanized. Primary tumors and lungs were collected for downstream analyses. Tumor tissue was divided and either snapfrozen for protein and DNA analysis, preserved in RNAlater for RNA analysis, or fixed in 4% paraformaldehyde for paraffin embedding.

#### Pulmonary metastasis quantification using Indian ink

Pulmonary metastatic burden was assessed by Indian ink inflation of the lungs. Following euthanasia, the trachea was exposed and a 15% solution of Indian ink in DEPC-treated water was injected intratracheally using a 5-mL syringe fitted with a 20-G needle until the lungs were completely inflated. Lungs were excised, briefly rinsed with water, and transferred to Fekete’s solution containing 90% 70% ethanol, 9% 37% formaldehyde, and 1% glacial acetic acid. Samples were stored at room temperature until analysis.

Lungs were visualized using a Nikon macroscope. Metastatic nodules appeared as white lesions against the black-stained lung tissue and were quantified manually.

#### *In vivo* GCN2-IN-6 treatment of solid cancer PDX and CDX models

Breast cancer patient derived xenograft (PDX) models for the experiments were obtained either from Jackson Laboratory (TM00098, TM00096), from TRACE PDX repository (BRC0004), or generated in-house (TNBC PDX 4) and passaged in female NSG mice. At the start of the experiment, tumor cell suspension of the PDX was implanted subcutaneously with Matrigel in the flank of female NSG mice (or NMRI-Foxn1^nu^ nude mice in the case of BRC0004) and once the tumors were measurable, mice were randomly divided into treatment groups (n = 5 per group for TM00098, TM00096 and TNBC PDX 4, n = 7 for BRC0004). GCN2-IN-6 was administered orally twice weekly at a concentration of 3 mg/kg. Mice were sacrificed once the first mouse reached a tumor volume above 2000 mm³ and primary tumors were harvested and processed identically as mentioned previously. Monotherapy of GCN2-IN-6 in TM00098, TM00096 and TNBC PDX 4 was tested at Ghent University, while for BRC0004, testing was performed at KULeuven TRACE PDX repository.

A total of 1.5 million A2780 ovarian cells were injected subcutaneously into the flank of 6-week-old female NSG mice to establish cell line-derived xenografts. Cells were suspended in a vehicle/Matrigel mixture at 30:70 ratio prior to injection. Once tumors were 150mm³, mice were randomized to receive vehicle or GCN2-IN-6 (3 mg/kg orally twice weekly; n = 6/group). Tumor dimensions were measured at least twice per week TM900v2 Tumor Volume Measurement System (BuDeTec). Animals were euthanized at 1500 mm³.

The neuroblastoma NEC005 PDX model was obtained through collaboration with the Experimental Pharmacology and Oncology (EPO) lab (Berlin, Germany) and expanded at Ghent University Hospital through subcutaneous injection into NSG mice. For subcutaneous injections, 100μL of mechanically dissociated tumor with 30-50% Matrigel (Corning, Cat # 356234) was injected subcutaneously with a 13G needle into the flank of the mice. Once tumors reached 150 mm³, mice were randomized to receive vehicle or GCN2-IN-6 (3 mg/kg orally twice weekly; n = 4/group). Tumor dimensions were measured at least twice per week using the TM900v2 Tumor Volume Measurement System (BuDeTec). Tumors were harvested when the control cohort reached the predefined endpoint of 1200 mm³.

#### *In vivo* CRISPRi experiments

MDAMB231 gSCRLi, gGCN2i #1 and gGCN2i #2 cells were harvested and resuspended in PBS. GCN2 knockdown was confirmed by RT-qPCR and western blot prior to the start of the experiment. A total of 3 × 10⁶ cells were injected subcutaneously into the flank of female NSG mice. Per cell line, once the tumors were measurable (volume > 62.5 mm³), the mice were randomized over two treatment groups (n = 5/group). In the first group CRISPRi-mediated knockdown was induced by continuous administration of doxycycline-infused chow (200 ppm). The second group received control chow. Tumor dimensions were measured at least twice per week using the TM900v2 Tumor Volume Measurement System (BuDeTec).

When the first animal reached a primary tumor volume > 2000 mm³, all mice were euthanized. Primary tumors were collected for downstream analyses. Tumor tissue was divided and either snapfrozen for protein and DNA analysis, preserved in RNAlater for RNA analysis, or fixed in 4% paraformaldehyde for paraffin embedding.

#### Combined ribosome profiling and RNAseq analysis wetlab

Primary tumors were collected from mice, snapfrozen in liquid nitrogen and stored at -80°C until further processing. Tissues were pulverized with a cell crusher and liquid nitrogen. The resulting tissue powder was resuspended with 750µl of lysis buffer supplemented with 100 mg/ml cycloheximide on ice. Efficient cell lysis was accomplished by passing the suspension through a 18G needle attached to a syringe. The cell suspension was incubated in lysis buffer for 10 min and centrifuged twice for complete removal of the cell debris. The clear lysate was aliquoted for subsequent footprinting, and an aliquot was used for total RNA purification. The RNA concentration of the lysates was determined with the Ǫubit RNA HS assay kit.

Total RNA from 2 µg of the lysate was purified with the RNA Clean and Concentrator-5 kit (Zymo Research). The profile of the eluted total RNA was visualized on Bioanalyzer with the RNA 6000 Nano kit for quality control. 10 µg of lysate was digested with RNase I (Thermofisher) at 21°C for 45 min. The reaction was stopped with the addition of Superase In RNase Inhibitor (Thermofisher). The digested lysate was purified on a MicroSpin S-400 HR Sephacryl column (GE Healthcare). The flow through was collected and purified with the RNA Clean and Concentrator-5 (ZymoResearch). The footprints were visualized on the Agilent Bioanalyzer with the RNA 6000 Nano kit (Agilent). Ribosome-protected fragments (RPFs) of 28-30 nt were selected by loading the footprints on a Novex TBE-Urea gel (15%) (Thermofisher) and performing gel electrophoresis. The gel parts corresponding to the fragments of interest were excised and left to diffuse in molecular grade water supplemented with ammonium acetate and SDS overnight at 4°C. The slurry was then transferred to CoStar filter tubes and centrifuged. The flow through was incubated at -80°C for 2h with glycogen and isopropanol. Following centrifugation at 18000 x g for 20 min, the supernatant was removed and the pellet was washed with ice-cold 80% EtOH. Pellets were left to dry for 10 min and were resuspended in 15 µl molecular grade water.

Initially, the RPFs were 3’ dephosphorylated with T4 PNK (New England Biolabs) for 1 h at 37°C to heal the 2’-3’ cyclic phosphate from the RNase I cleavage and were subsequently purified using the Oligo Clean and Concentrator kit (ZymoResearch). The 3’ adapter was ligated to the RPFs and the ligated fragments were purified following the protocol of the Small RNA Library prep kit (Lexogen). After the 5’ phosphorylation of the RPFs and their purification with the Oligo Clean and Concentrator kit, 5’ adapter ligation took place based on the Small RNA Library prep kit. Ribosomal RNA (rRNA) was depleted with the addition of LNA probes from the ǪIAseq FastSelect rRNA HMR kit (Ǫiagen) during the cDNA synthesis reaction. Library products were purified with AMPure XP beads and their quality was determined on the Agilent Bioanalyzer with the DNA High Sensitivity kit (Agilent). For the efficient removal of adapter dimers and excess index primers, the library products run on a E-Gel EX Agarose gel (4%) (Thermofisher) and the gel part corresponding to the library products was excised and purified.

400 ng of total RNA material from Step 2.3 was used further for matching mRNA-seq. mRNA was first enriched using oligo(dT) beads and was fragmented randomly by adding fragmentation buffer. Then, cDNA was synthesized by using random primers, after which a second-strand synthesis buffer, dNTPs, RNase H and DNA polymerase I were added to initiate the second-strand synthesis. Following end repair and sequencing adaptor ligation, the double-stranded cDNA library was completed through size selection and PCR enrichment. The concentration of the libraries was estimated by performing qPCR using a dilution of PhiX Control v3 as standard and samples were pooled in an equimolar way. High coverage sequencing was performed on the Illumina Nextseq 500 system with the high output v2 kit and SE75 configuration.

#### Reference information RIBOseq and RNAseq

The necessary reference information was obtained from ftp.ensembl.org. For genome annotation information, Ensembl releases 105 was used for the Homo sapiens (human) data (assembly GRCh38). Human gene annotation information in GTF format and reference sequences in FASTA format were also obtained from ftp.ensemblgenomes.org. Additional reference information for prefiltering purposes includes: (i) human rRNA nucleotide sequences from ncbi.nlm.nih.gov/nuccore (obtained using the search term ‘biomol_rrna’), (ii) human sn-RNA and sn-o-RNA sequences from Biomart (obtained using the gene type filters for ‘sn-RNA’ and ‘sn-o-RNA’) and (iii) human tRNA sequences from gtrnadb.ucsc.edu.

#### Drylab RIBOseq

Raw ribosome profiling sequences in FASTǪ format were checked for general quality with FastǪC, executed under the default settings suggested by the FastǪC program. Adaptor sequence ‘TGGAATTCTCGGGTGCCAAGG’ was clipped and sequences were trimmed based on sequencing quality in all datasets with the Trimmomatic toolset (0.39). Sequences shorter than 20 nucleotides were hereby discarded and the quality threshold was set at 28. Afterwards, all contaminant PhiX, rRNA, snRNA, sn-o-RNA and tRNA sequences were filtered before mapping the remaining reads to the human genome. For all these filtering and mapping steps, the STAR (version 2.7.2b) splice-aware mapping option was set. Only uniquely mapped reads were considered. A tabular output (*Table S4)* is provided with the mapping statistics (prefiltering against contaminant rRNA, snRNA, sn-o-RNA and tRNA sequences and mapping the remaining reads to the human genome). FastǪC was executed a second time under default settings on the aligned sequences in the SAM files to assess the quality of the ribosome protected fragments (RPFs). As extra quality control the offset between the 5’ site of the ribosome protected fragment (RPF) and the P-site was calculated for each RPF length with the tool Plastid. Differential translation, transcription and translation efficiency analysis was performed using DESeq2 (version 1.42.1) and the deltaTE method^1^.

#### RNA sequencing of CRISPRi and GCN2-IN-6 treated samples

Multiplexed libraries from isolated RNA were prepared using the ǪuantSeq 3’ mRNA FWD kit (Lexogen, Vienna, Austria) and sequenced on an AVITI24 to obtain single-end reads with a length of 100 nucleotides to a target depth of 30M reads per sample. Reads were trimmed with TrimGalore (version 0.6.10) and mapped to human genome assembly GRCh38 and accompanying gtf file using STAR (version 2.7.11b). One sample (a sextuplet of doxycycline induced in vivo samples with gGCN2i3 construct) was excluded due to low library concentration with subsequent low read counts and few detected genes. Gene-level read counts provided by STAR were analyzed in R (version 4.5.1) using DESeq2 (version 1.48.2). Filtering criteria for genes were >= 100 reads in >=20 of the 37 retained samples leaving 9,946 genes for analysis. For gene-set enrichment analyses, genes were ranked using their signed DESeq2 Wald statistic.

#### Analysis METABRIC database

Expression log intensity levels from publicly available mRNA microarray expression data was obtained from cBioportal. The Z-score was calculated for expression levels of *ATF4.* To subdivide patients into an *ATF4^high^*and Other group, we selected samples with Z-score > 1. Calculation of differentially expressed genes between the two groups was assessed using limma (version 3.50.3). Using differentially expressed genes that had an absolute logFC > 0.5 and pval < 0.05, gene set enrichment analysis (core analysis) was performed.

